# Evolutionary insights into β-cyclocitral signaling in *Physcomitrium patens*

**DOI:** 10.64898/2026.10.13.758730

**Authors:** Paolo Korwin Krukowski, Chiara Toffanin, Maurizio Carnà, Edoardo Tosato, Claudia Beraldo, Alessandro Alboresi, Stefano D’Alessandro

## Abstract

- How did a product of photooxidative damage become a signal for stress tolerance? β-cyclocitral (β-cc) promotes acclimation in several angiosperms, yet has detrimental effects in other photosynthetic organisms. We investigated this contrast in the moss *Physcomitrium patens* to explore the evolution of β-cc signaling.
- We combined apocarotenoid treatments and excess light exposure with photosynthetic measurements, biochemical analyses and RNA sequencing. Comparative phylogenomics traced the conservation of regulatory and detoxification components associated with β-cc responses.
- Physcomitrium accumulated β-cc under excess light and mounted a strong transcriptional response to exogenous β-cc, substantially overlapping with its light-stress response. Nevertheless, β-cc impaired photosynthetic performance and provided no protection. At a phylogenetic level, while MBS1 was deeply conserved, the downstream detoxification module showed marked divergence. Notably, ANAC081/ANAC102-like proteins were recovered in terrestrial monocots but not in Physcomitrium or *Lemna turionifera*, both showing detrimental responses to β-cc.
- Our findings suggest that the evolution of protective β-cc signaling involved the recruitment of downstream detoxification regulators, with the SCL14/TGAII- and ANAC081/ANAC102-related lineage emerging as a candidate link between an ancient oxidative cue and beneficial acclimation.

## 1 Introduction

Carotenoids are pigments that occur in Archaea and Bacteria and pre-date the emergence of oxygenic photosynthesis (Britton *et al*., 2012; Sandmann, 2021). Although largely hydrophobic, many contain polar functional groups at one or both ends of the molecule, with lycopene and β-carotene being notable exceptions. These structural properties influence their positioning within membranes and pigment–protein complexes (Bassi *et al*., 1993; Gruszecki & Strzałka, 2005; Dall’Osto *et al*., 2015). Following the emergence of photosynthetic eukaryotes, carotenoid biosynthesis diversified across algal lineages, expanding the range of pigments involved in light harvesting and photoprotection. The pathway underwent further modifications and in green algae and land plants includes the hydroxylation of α-carotene to produce lutein and the epoxidation of zeaxanthin to violaxanthin, which is subsequently converted into neoxanthin (Sandmann, 2021).

β-carotene is an integral component of the photosynthetic reaction-center core complexes and plays a central role in photoprotection, a function conserved since the emergence of cyanobacterial photosynthesis (Telfer, 2002; Ramel *et al*., 2012; Pinnola & Bassi, 2018). Oxidative cleavage of carotenoids generates smaller compounds collectively known as apocarotenoids (Stratton *et al*., 1993; Sommerburg *et al*., 2003; Ramel *et al*., 2012; Burton *et al*., 2021). The best-characterized apocarotenoids are those that function as plant hormones, such as abscisic acid (ABA), strigolactones (SLs) and zaxinone, as well as other apocarotenoids that are emerging as signaling molecules (Wang *et al*., 2019; Simkin, 2021; Jia *et al*., 2022; Carnà *et al*., 2025).

β-cyclocitral (β-cc) is a bioactive volatile apocarotenoid generated by the oxidative cleavage of β-carotene (Carnà *et al*., 2025). Along with β-ionone and dihydroactinidiolide, β-cc has been reported to act as a signaling molecule enhancing tolerance to abiotic and biotic stress (Shumbe *et al*., 2014; Felemban *et al*., 2024). Notably, the physiological effects of β-cc vary considerably among photosynthetic lineages and species. β-cc can exert toxic effects on cyanobacteria and algae whereas it is predominantly associated with enhanced stress tolerance in terrestrial plants (Tomita *et al*., 2016; D’Alessandro *et al*., 2018; Roach *et al*., 2022; Wang *et al*., 2024). In cyanobacteria and some algae, β-cc has been reported to repress chlorophyll biosynthesis. In the green alga *Chlamydomonas reinhardtii*, exposure to β-cc vapor decreased the maximum quantum yield of photosystem II (PSII) and reduced non-photochemical quenching (NPQ) (Roach *et al*., 2022).

Strong inhibitory effects, including the activation of cell-death processes, have also been reported in the cyanobacterium *Microcystis aeruginosa*, the green alga *C. reinhardtii* and the aquatic angiosperm *Lemna turionifera* (Sun *et al*., 2020; Pan *et al*., 2024; Wang *et al*., 2024). In terrestrial plants, however, β-cc is predominantly described as a product of β-carotene photooxidation under conditions of excessive PSII excitation. It acts as a signaling molecule that promotes acclimation and enhances tolerance to environmental stress (Ramel *et al*., 2012; Lv *et al*., 2015; Mitra *et al*., 2021; Avesani *et al*., 2025). Exogenous β-cc treatments elicit extensive transcriptional and metabolic reprogramming in several angiosperms, including *Arabidopsis thaliana*, *Solanum lycopersicum*, *Oryza sativa* and *Vitis vinifera* (Deshpande *et al*., 2021; Taniguchi *et al*., 2023; Tiwari *et al*., 2025).

Mechanistically, β-cc-induced tolerance to excess light (EL) has been associated with METHYLENE BLUE SENSITIVITY 1 (MBS1)–dependent regulation of singlet-oxygen–responsive genes (SORGs) and the activation of the SCARECROW-LIKE 14 (SCL14)–dependent xenobiotic detoxification pathway (Shumbe *et al*., 2017; D’Alessandro *et al*., 2018). β-cc has also been reported to interact with phytohormone signaling (Lv *et al*., 2015; Taniguchi *et al*., 2023) and shown to inhibit 1-deoxy-D-xylulose 5-phosphate synthase (DXS), although only at concentrations observed exclusively in cyanobacteria so far (Jüttner *et al*., 2010; Mitra *et al*., 2021).

Despite these advances, how β-cc responses vary across the green lineage and which downstream pathways underpin its beneficial effects in terrestrial plants remain poorly understood. Here, we address these questions by investigating β-cc responses in *Physcomitrium patens*, a bryophyte model belonging to an early-diverging land-plant lineage. It retains several ancestral traits and shares features of its photosynthetic machinery with both algae and vascular plants (Beraldo *et al*., 2025), making it particularly suitable for studying β-cc responses and the evolution of retrograde signaling.

## 2 Materials and Methods

### 2.1 Plant growth conditions and treatments

Protonemal tissue of *Physcomitrium patens* was grown on rich medium (PPNH_4_) (Ashton *et al*., 1979) for 10 days in 10-cm Petri dishes under 60 µmol m^−2^ s^−1^ 4000K white light, with a 12-h light/12-h dark photoperiod and day/night temperatures of 24°C/20°C. Protonemal tissue was then harvested and transferred in spots to new plates containing minimal medium (PPNO_3_), where it was grown for a further 18 days (Ashton *et al*., 1979). All experiments were performed on plates containing 6-10 spots of Physcomitrium.

Four-week-old plants were exposed to excess light stress by transferring them to 1500 µmol m^−2^ s^−1^ photosynthetic photon flux density (PPFD) of 4000K white light and constant airflow at room temperature (RT).

β-cc (90% purity) Thermo Scientific (B25700.09, lot 10237632). β-I (96% purity) Thermo Scientific (297131000, lot A0461319). β-cc and β-I pretreatments were performed by exposing plants to volatile β-cc or β-I for 4 h in a transparent 10-L desiccator under light (60 μmol photons m^−2^ s^−1^) at RT. An aliquot of β-cc was deposited on a cotton wick to promote volatilization within the airtight desiccator. For control treatments, β-cc was replaced with distilled water.

### 2.2 PAM Imaging and data analysis

4-week-old plants were dark-adapted for 20 min, and chlorophyll fluorescence was measured using a Fluorcam FC 800-O/1010 imaging fluorometer (Photon System Instruments). Chlorophyll fluorescence was recorded using a modified light curve protocol, in which plants were exposed to successive 1-min steps of actinic white light at PPFDs of 0.82, 56.05, 92.87, 148.1, 240.15 and 332.2 μmol photons m^−2^ s^−1^. F_0_ was measured immediately after dark adaptation and a subsequent 6s exposure to far-red light. Far-red light was also applied after each actinic light step to measure F_0_’. F_m_ was measured using a saturating white-light pulse at 2500 μmol photons m^−2^ s^−1^. The maximum quantum yield of PSII (QY_max_) was calculated as (F_m_ – F_0_)/F_m_. NPQ_(T)_ was calculated as the mean (F_m_/F_0_-1) of control conditions divided by the (F_m_’/F_0_’-1) of the sample −1 (Tietz *et al*., 2017). qP was calculated as F_q_’/F_v_’ with F_q_’ = F_m_’-F_t_’, where F_t_’ is the chlorophyll fluorescence measured at the end of each actinic light period, before the saturating pulse and far-red illumination.

### 2.3 GC-MS quantification of apocarotenoids

4-week-old plants were exposed to EL conditions. Pools of two Petri dishes (10 spots) were sampled and immediately frozen in liquid nitrogen. Samples were ground to a powder, and 200 mg of powder was transferred to an extraction buffer containing 1 mL of 1mM HCl, 2 mL of methyl *tert*-butyl ether (MTBE) and 100 ng mL^−1^ linalool (97% purity) as reference (Sigma Aldrich). Samples were vortexed and sonicated for 10 min in ice-cold water. Following centrifugation (5 min at 5000 g) the supernatant was transferred to a fresh container and concentrated under a stream of nitrogen while kept on ice.

Analyses were performed using an Agilent 6890N gas chromatograph (GC) coupled to an Agilent 5973 mass selective detector (MS) and fitted with a CPS CC-5MS-Xil Column. Authentic standards were analyzed to determine retention times and confirm mass spectra. Selected-ion monitoring (SIM) was performed using characteristic fragment ions at m/z 137 for β-cc, m/z 153 for β-cca, m/z 177 for β-I and m/z 93 for linalool. Helium was used as the carrier gas at a flow rate of 1 mL min^−1^. The oven temperature was initially set to 60°C and increased 3°C min^−1^ to 150°C, followed by an 8-min column-cleaning step at 300°C.

The area of each analyte was normalized to that of linalool.

### 2.4 MDA quantification

4-week-old plants pretreated with β-cc or water were exposed to EL conditions. Two Petri dishes (10 spots) were sampled and immediately frozen in liquid nitrogen. Samples were ground to a powder, and 100 mg of powder was used for extraction. 1 mL of extraction buffer (80% Ethanol, 0.01% butylated hydroxytoluene) was added to the powder. Samples were sonicated for 3 min in ice-cold water and centrifuged at 4°C for 10 min at 11000g. Two 400-µl aliquots of the supernatant were transferred to separate centrifuge tubes, and 1.5 volumes of reaction buffer (20% trichloroacetic acid with or without 0.065 g mL^−1^ thiobarbituric acid) were added to each aliquot. Samples were incubated 10 min at 93°C, and absorbance was measured at 532 nm and 600 nm. MDA was quantified using a calibration curve prepared with tetramethoxypropane (Angene) and the procedure described by (Hodges *et al*., 1999; Landi, 2017).

### 2.5 Fatty acid methyl esters analysis

4-week-old plants were exposed to β-cc or water. Two Petri dishes (10 spots) were sampled and immediately frozen in liquid nitrogen. Samples were ground to a powder, and 100 mg of powder was subjected to acid-catalyzed methylation using boron trifluoride (10% w/v in methanol) to obtain fatty acid methyl esters (FAME) (Christie & Han, 2010). FAMEs were identified by GC-MS and quantified by GC-FID (GC-2010 Plus, Shimadzu, Japan). Helium was used as the carrier gas at a constant flow rate of 1 mL min^−1^, and separation was achieved using a nonpolar ZB5-MS capillary column (Phenomenex, USA). The injector temperature was set to 250°C. The oven temperature was initially held at 60°C for 1 min, increased to 180°C at 10°C min^−1^ and held for 1 min, then increased to 230°C at 1°C min-1 and held for 2 min. Finally, the temperature was increased to 320°C at 15°C min-1 and held for 5 min. The same column and chromatographic conditions were used for GC-MS and GC-FID analyses. Compounds were identified by comparing their mass fragmentation spectra and retention indices with reference values (Haslam *et al*., 2024).

### 2.6 Protein extraction and immunodetection

Total protein extracts from samples exposed to β-cc for 4h or left untreated were prepared by grinding tissues in sample buffer (50 mM Tris pH 6.8, 100 mM DTT, 2% SDS, 10% glycerol) and resolved by SDS–PAGE. Sample volumes corresponding to equivalent amounts of chlorophyll were loaded. Following electrophoresis, proteins were transferred onto nitrocellulose membranes and detected using in-house polyclonal primary antibodies followed by an alkaline phosphatase–conjugated secondary antibody (Sigma, #A3562).

For densitometric analysis of bands, images were processed in Fiji (https://fiji.sc/). Gray values were measured for each band, and a local background was subtracted. Signal intensities were then normalized to the untreated wild-type (WT).

### 2.7 RNA sequencing and analyses

4-week-old plants were treated with β-cc for 4 h or exposed to EL conditions for 16 h. Three pools, each comprising two Petri dishes (10 spots) were sampled and immediately frozen in liquid nitrogen. Samples were ground to a powder, and RNA was extracted from 100 mg of powder using TriFast (30-2010, VWR) following the manufacturer’s instructions. RNA-seq libraries were prepared and sequenced by the DiBio sequencing service (<u>University</u> of Padova). Sequencing was performed in paired-end mode, targeting 50 million reads per sample. Adapter trimming and read quality filtering were performed by the sequencing service before downstream analyses.

Transcript abundance was quantified using Salmon with indexes built from the *P. patens* transcriptome (Ppatens_947_v7.1.transcript and Ppatens_318_v3.3.transcript separately) using the genomic sequence as a decoy (Goodstein *et al*., 2012; Bi *et al*., 2024). Counts were imported into R and summarized at the gene level for differential expression analysis, accounting for transcript length and library size effects in downstream modeling (Soneson *et al*., 2016). Differential expression was tested in R using DESeq2 (Love *et al*., 2014). Gene-level count matrices were modeled using a negative binomial generalized linear model, with the default size-factor normalization and dispersion estimation procedures, implemented in DESeq2. Differential expression was assessed using Wald tests. P values were adjusted for multiple testing using the Benjamini–Hochberg method to control the false discovery rate (FDR). Genes were considered differentially expressed if they met the following criteria: a minimum of 10 reads per sample, an adjusted p-value < 0.01 and a log_2_ fold change (LFC) < −1 or > 1 (Supplemental Data 1). To obtain stabilized effect sizes, LFCs were shrunk using the ashr method. Shrunken LFCs were used for fold-change reporting and ranking (Stephens, 2017).

Downstream analyses and visualization were performed in R using tidyverse-compatible workflows (readr, dplyr, tidyr, purrr, fs) and the base R stats package. Plots were generated using ggplot2, with label placement optimized via ggrepel (Wickham, 2016). Sample-to-sample relationships and expression patterns were visualized using heatmaps generated with pheatmap. Overlaps among gene sets were visualized using VennDiagram. Functional enrichment analyses were performed using clusterProfiler, with org.At.tair.db and AnnotationDbi providing Arabidopsis gene annotations based on a published homology analysis (Yu, 2024). Enrichment and functional profiling were also corroborated using gprofiler2. Where needed, metadata and supplementary tables were imported using readxl. Parallel computation for selected steps was enabled via BiocParallel.

A rank-overlap enrichment was assessed separately in *Arabidopsis thaliana*, adapted from (Carna *et al*., 2026) and *Physcomitrium patens*. Genes with adjusted p<0.01 in at least one contrast were ranked by fold change. Concordant and discordant overlaps were tested across each ranking using one-sided hypergeometric tests. P-values were Bonferroni-corrected across the two directional overlaps and adjusted using the Benjamini–Yekutieli method across 100 tests per species. Adjusted p<0.01 was considered significant. Spearman correlations were calculated for the selected genes.

### 2.8 Comparative proteome and phylogenetic analyses

A comparative proteomic dataset was assembled to cover major lineages of photosynthetic organisms. The dataset comprised 14 proteomes: *Arabidopsis thaliana* (TAIR12), *Arabidopsis lyrata* (v1.0), *Solanum lycopersicum* (SL4.0), *Hordeum vulgare cv. Morex* (MorexV3), *Lemna turionifera 9434* (CSHL2022v1), *Amborella trichopoda* (AMTR1.0), *Picea abies* (v1.0b), *Selaginella moellendorffii* (v1.0), *Physcomitrium patens* (v7), *Marchantia polymorpha* (v7.1), *Mesotaenium endlicherianum* (v2), *Zygnema circumcarinatum* SAG 698-1b, *Chlamydomonas reinhardtii* (v5.5), and *Microcystis aeruginosa* NIES-843. For Figure S8 an expanded dataset was analyzed adding: *Brachypodium distachyon* (v3.0), *Brassica rapa* (Brapa1.0), *Eutrema salsugineum* (Eutsalg1_0), *Glycine max* (v2.1), *Oryza sativa* (IRGSP-1.0), *Populus trichocarpa* (Pop_tri_v4), *Spirodela polyrhiza* (v2.290), *Triticum aestivum* (IWGSC), *Vitis vinifera* (ASM3070453v1) and *Zostera marina* (Zosma v2.1). Proteomes were standardized to reduce biases, and a single representative protein per gene was retained by selecting the longest translated isoform. Following preprocessing, the combined dataset contained 344,698 protein sequences. Orthogroups were inferred using OrthoFinder v2.5.5 (Emms & Kelly, 2019). All-versus-all protein similarity searches were performed with DIAMOND. Orthogroups relevant to β-cc signaling and detoxification were identified using experimentally characterized *A. thaliana* proteins as references. The NAC transcription factors ANAC002 (*AT1G01720*), ANAC032 (*AT1G77450*), ANAC081 (*AT5G08790*), and ANAC102 (*AT5G63790*) were used to identify the NAC orthogroup containing the β-cc-associated ATAF proteins. SCL14 (*AT1G07530*) was used to identify the corresponding GRAS-family orthogroup, whereas MBS1 (*AT3G02790*) was used as the reference for the MBS1 orthogroup. The class II TGA transcription factors TGA2 (*AT5G06950*), TGA5 (*AT5G06960*), and TGA6 (*AT3G12250*) were used to identify the TGA-containing orthogroup. Detoxification-related reference proteins included CYP81D11 (*AT3G28740*), AER (*AT5G16970*), the aldo-keto reductases AKR4C8 (*AT2G37760*) and AKR4C9 (*AT2G37770*), the chloroplast aldehyde reductases *AT3G04000* and *AT1G54870*, SDR1 (*AT3G61220*), and selected glutathione S-transferases. Protein sequences belonging to each target orthogroup were extracted from the OrthoFinder results and aligned with MAFFT v7.505 (Katoh & Standley, 2013). Maximum-likelihood phylogenetic trees were reconstructed from the protein alignments using IQ-TREE v2.0.7 (Minh *et al*., 2020). The optimal amino-acid substitution model was selected independently for each alignment using ModelFinder according to the Bayesian information criterion (Kalyaanamoorthy *et al*., 2017). Branch support was evaluated using 1,000 ultrafast bootstrap replicates and 1,000 SH-like approximate likelihood-ratio test (SH-aLRT) replicates. For graphical presentation, internal-node support values were displayed when SH-aLRT was ≥70 and/or ultrafast bootstrap support was ≥75; both statistics are reported as SH-aLRT/UFBoot. Phylogenetic placement, rather than orthogroup membership alone, was used to assess conservation of the specific β-cc-associated protein lineages.

Pairwise amino-acid identity and sequence coverage were calculated between *A. thaliana* reference proteins and candidate homologs. Identity was calculated over aligned positions containing amino acids in both sequences, whereas coverage was expressed as the proportion of residues in the *A. thaliana* reference sequence aligned to a residue in the compared sequence.

### 2.9 *Cis*-regulatory motif enrichment analysis

Promoter regions spanning −1000 to +100 bp relative to the annotated transcription start site were extracted for β-cc-upregulated and background genes. Motif occurrences were identified with FIMO using p<0.0001 (Grant *et al*., 2011) with the maximum number of stored scores increased to 1000000. JASPAR plant motifs were obtained from the JASPAR 2024 CORE collection (Rauluseviciute *et al*., 2024) and grouped into transcription factor families for visualization. Enrichment was assessed using two-sided Fisher’s exact tests with Benjamini–Hochberg correction. Motifs with FDR < 0.05 were considered significant. For strongly induced genes, genes with log2FC ≥ 1.5 were compared with the remaining β-cc-upregulated genes and motifs with FDR < 0.05 were retained.

De novo motif discovery was performed using STREME (Bailey, 2021). The as-1/TGA-like motif (TGACGTCAKBRMTKACGT) (Fode *et al*., 2008) and NAC-related motifs were analyzed separately. NAC-any comprised ANAC102-like (CACGBMACY), ANAC032-like (CACGTAAC) (Song *et al*., 2016) and ANAC002-like (MA2015.2) motifs. Motif frequencies were compared between *A. thaliana* and *P. patens* among the top 100, 200 and 500 β-cc-upregulated genes and among genes with log_2_FC ≥ 2 or ≥ 3 using Fisher’s exact tests followed by Benjamini–Hochberg correction.

### 2.10 Data analysis and availability

Graphs were generated in Python (Python Software Foundation, https://python.org/) using the Pandas, Matplotlib, and Seaborn libraries (Hunter, 2007; McKinney, 2010; Waskom, 2021).

Statistical tests were performed using the Pingouin and SciPy libraries (Vallat, 2018; Virtanen *et al*., 2020). The Games-Howell post hoc test was used for multiple comparisons of normally distributed data, and the Mann-Whitney U test was used for comparisons between independent groups when a nonparametric approach was required. All experiments were performed in at least two complete experimental replicates, each comprising at least two Petri plates per condition. Each Physcomitrium spot has been treated as an independent sample.

Scripts are available in the supplementary data files.

## 3 Results

### 3.1 β-cc alters photosynthetic performance in Physcomitrium

We assessed photosynthetic performance in β-cc-pretreated and control plants using pulsed-amplitude modulation (PAM) imaging. Three 10 cm round Petri dishes, each containing 6 to 8 spots of 4-week-old plants, were exposed to control conditions or to β-cc vapor generated from increasing volumes of the compound (10, 25, 50 or 100 μL) at RT for 4 h. As reported for some aquatic photosynthetic organisms, Physcomitrium showed a reduction in the maximum quantum yield of PSII (QY_max_, Figure 1a, b) and in NPQ induction (Figure 1c) (Mitra *et al*., 2021; Roach *et al*., 2022). Specifically, plants under control conditions showed a QY_max_ of 0.75, while increasing β-cc treatment volumes led to QY_max_ values of 0.73, 0.71, 0.70 and 0.68, respectively. Similarly, β-cc treatments with volumes greater than 10 μL reduced NPQ induction, especially at high PPFD (250-350 µE, Figure 1c). These results were unexpected, as β-cc treatments have been reported to leave QY_max_ unaffected and to exert beneficial priming effects on photosynthesis in other land plants.

**Figure 1:**
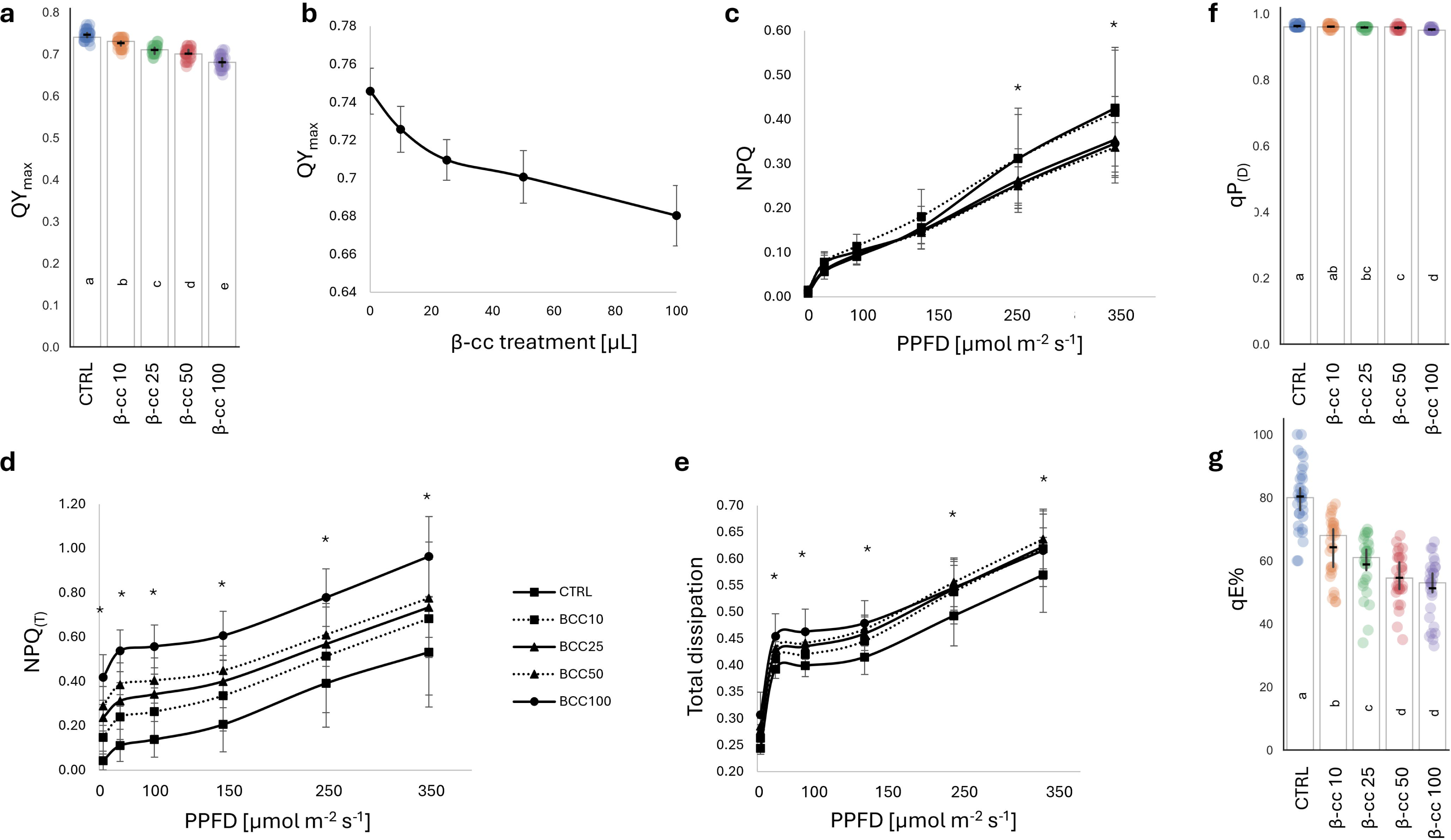
β-cyclocitral reduces photosynthetic performance and increases sustained energy dissipation in Physcomitrium. 28-day after inoculation Physcomitrium plants were exposed for 4 h to volatile β-cyclocitral (β-cc; 10, 25, 50 or 100 µL) or to control conditions, and chlorophyll fluorescence parameters were measured by PAM imaging. (a) Maximum quantum yield of PSII (QY_max_) after β-cc treatment. (b) Dose dependence of QY_max_. (c) NPQ induction during the light-response curve. (d) NPQ_(T)_, calculated relative to control values. (e) Total energy dissipation, calculated as ΨNPQ + ΨNO. (f) Photochemical quenching after 1 min of darkness following the light curve, qP_(D)_. (g) Rapidly reversible NPQ fraction after 1 min of darkness, qE%. Data are means ± SD. Asterisks indicate significant differences relative to the corresponding control. Alternatively, different letters are assigned to statistically different groups.

Using NPQ_(T)_, calculated relative to control values, we observed an increase in non-photochemical quenching with increasing β-cc treatment volumes (Figure 1d)(Tietz *et al*., 2017). β-cc increased NPQ_(T)_ in a dose-dependent manner, from 0.11 in control plants to 0.54 at the highest treatment volume under growth light (Fig. 1d). Consistently, total energy dissipation (Ψ_NPQ_ + Ψ_NO_) increased across light intensities, reaching 64% compared with 57% in controls at high irradiance (Figure 1e).

We also analyzed the recovery of chlorophyll fluorescence after 1 min of darkness at the end of the PAM measurements. This measurement allowed us to assess PSII reaction-center reopening and estimate photochemical quenching in the dark (qP_(D)_), which reflects the redox state of the PSII acceptor side, and the rapidly reversible fraction of NPQ (qE%). β-cc treatments only marginally affected the rapid reopening of PSII reaction centers in the dark (qP_(D)_ after 1 min of darkness, Figure 1f). By contrast, the rapidly reversible fraction of NPQ after 1 min of darkness (qE%) declined from approximately 80% in controls to 51% at the highest β-cc dose (Figure 1g), indicating an increased contribution of sustained quenching.

The negative effect of β-cc treatments on Physcomitrium photosynthetic performance resembles the response observed in aquatic organisms rather than the priming effect observed in other land plants and appears relatively insensitive to light intensity (Roach *et al*., 2022). We further investigated whether the β-cc-induced reduction in photosynthetic performance required light by performing a 50µl β-cc treatment under light and dark conditions. We chose this treatment volume as it is commonly used in the literature (D’Alessandro *et al*., 2018). Untreated Physcomitrium samples showed similar QY_max_ values of approximately 0.75 after 4 h under either light or dark conditions (Supplemental Figure S1a). The 50µl β-cc treatment decreased QY_max_ to 0.70 under light conditions and to a slightly lower value in darkness (0.69, Supplemental Figure S1a). Similarly, β-cc treatments under light and dark conditions produced broadly similar responses in NPQ_(T)_, total dissipation, qP_(D)_ and qE%, suggesting that the effects of β-cc on Physcomitrium do not require light during treatment (Supplemental Figure S1b, c, d, e).

**Figure S1:**
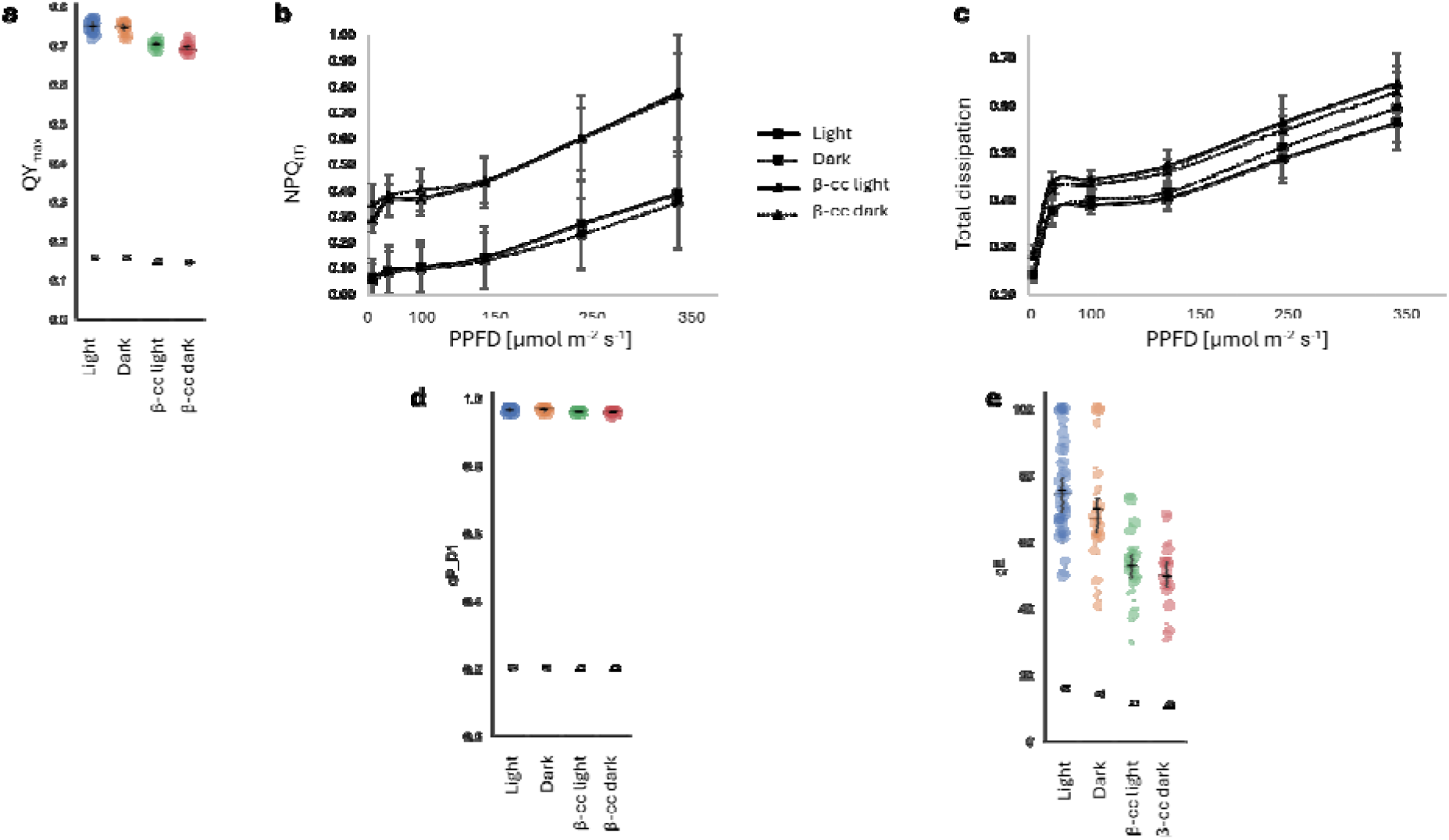
Effect of light during β-cyclocitral treatment on Physcomitrium photosynthetic parameters. 28-day after inoculation Physcomitrium plants were exposed for 4 h to volatile β-cc (50 µL) or control conditions either in the light or in darkness. (a) QY_max_. (b) NPQ_(T)_. (c) Total energy dissipation. (d) qP_(D)_ after 1 min of darkness. (e) qE% after 1 min of darkness. Data are means ± SD. Asterisks indicate significant differences relative to the corresponding control. Alternatively, different letters are assigned to statistically different groups.

### 3.2 β-cc reduces QY_max_ and photosynthetic performance under EL

We next tested whether β-cc could prime plants for exposure to excess light (EL, 1500 μmol photons m^−2^ s^−1^) conditions. Plates containing six to ten Physcomitrium spots were first treated with 50µl of β-cc and then exposed to EL conditions. Contrary to the protective response reported in other land plants, β-cc exacerbated EL-induced photoinhibition, reducing QYmax from 0.43 under EL alone to 0.27 in pretreated plants (Figure 2). Photoinhibition was more pronounced in the central regions of the spots reflecting the older age of the central tissue resulting from radial growth (Figure 2b) (Thelander *et al*., 2005; D’Alessandro *et al*., 2018). EL strongly increased NPQ_(T)_ and total energy dissipation with further increases following β-cc pretreatment (Figure 2c, d). Under growth light, approximately 40% of absorbed energy was dissipated, compared with 70% under EL and over 80% following β-cc pretreatment (Figure 2d). β-cc pretreatment also reduced qP, suggesting that the treatment affected the redox state of the PSII acceptor side (Figure 2e). While 1 min of darkness was sufficient to allow nearly complete reopening of PSII reaction centers under control conditions, the qP_(D)_ parameter resulted in only 60% of PSII reaction centers open after 4 h of EL and only 40% in β-cc-pretreated plants exposed to EL (Figure 2f).

**Figure 2:**
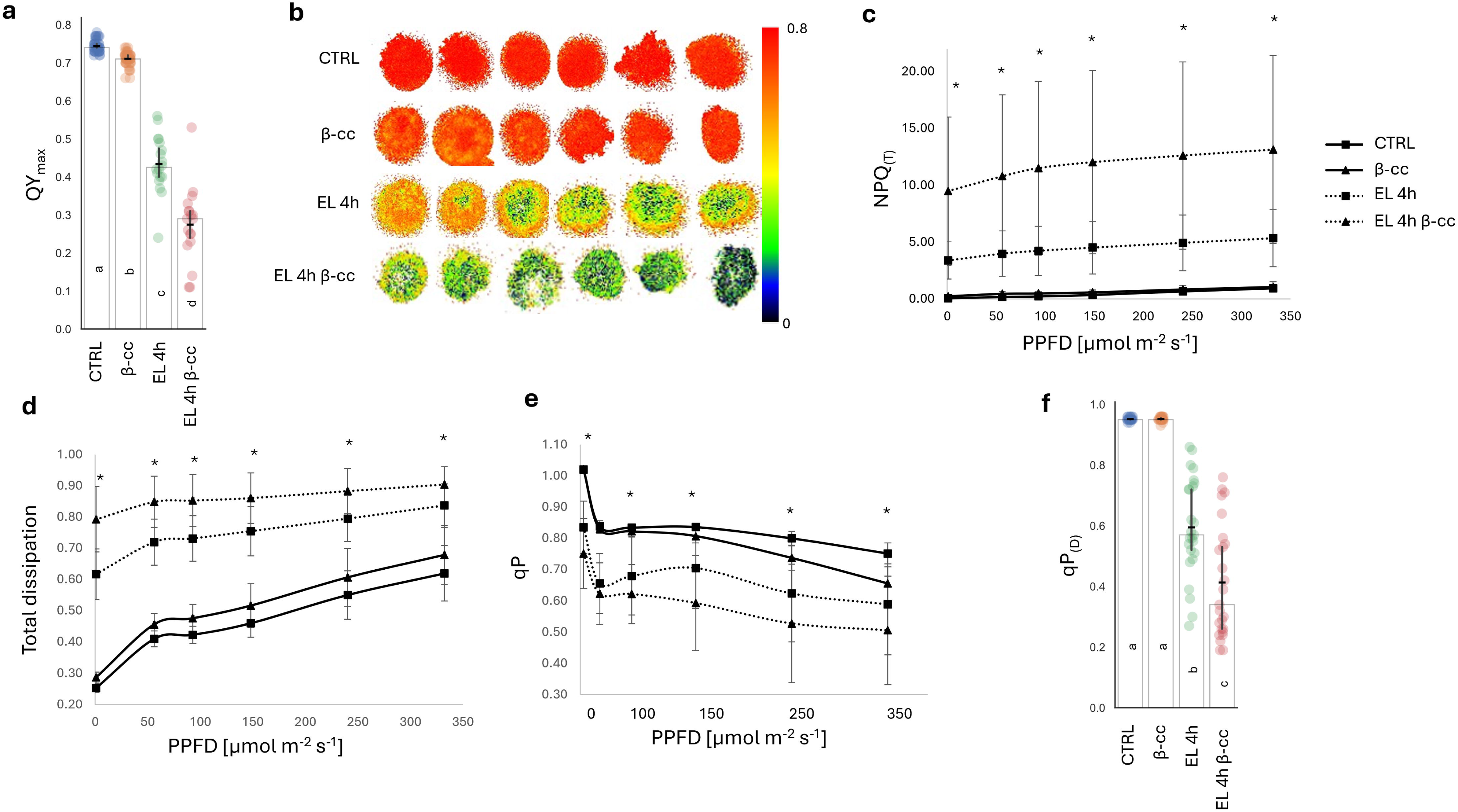
β-cyclocitral does not protect Physcomitrium from excess light and further impairs PSII performance. 28-day after inoculation Physcomitrium plants were pretreated for 4 h with volatile β-cc (50 µL) or control conditions and then exposed to excess light (EL; 1500 µmol photons m^−2^ s^−1^) for 4 h. (a) QY_max_ under control, β-cc, EL and EL + β-cc conditions. (b) Representative QY_max_ images showing spatial heterogeneity of EL-induced photoinhibition across spots. (c) NPQ_(T)_ during the light-response curve. (d) Total energy dissipation. (e) Photochemical quenching (qP) during the light-response curve. (f) qP_(D)_ after 1 min of darkness following the light curve. Data are means ± SD. Asterisks indicate significant differences relative to the corresponding control. Alternatively, different letters are assigned to statistically different groups.

### 3.3 Specificity of the photosynthetic response to β-cc

β-cc is only one of the apocarotenoids generated by β-carotene oxidation under EL (Ramel *et al*., 2012). β-cc can act as a retrograde signal, as can β-cyclogeranic acid (β-cyclocitric acid), dihydroactinidiolide and β-ionone (β-I) (Shumbe *et al*., 2014; D’Alessandro *et al*., 2019; Felemban *et al*., 2024). As the response to β-cc in Physcomitrium appeared detrimental rather than protective, we asked whether the observed effects reflected a response shared with other cyclic apocarotenoids or were specific to β-cc. To investigate this, we tested the effects of β-I in Physcomitrium by performing a 4 h pretreatment with vapor generated from 50 μL of β-I. Unlike β-cc, β-I did not reduce QY_max_(Figure 3a and b). Furthermore, NPQ_(T)_, total energy dissipation and qP did not differ significantly between β-I-treated and untreated plants under either control or EL conditions (Figure 3c, d, e, f). These results show that the detrimental effects of β-cc on photosynthetic performance were not reproduced by β-I, indicating a rather β-cc-specific response.

**Figure 3:**
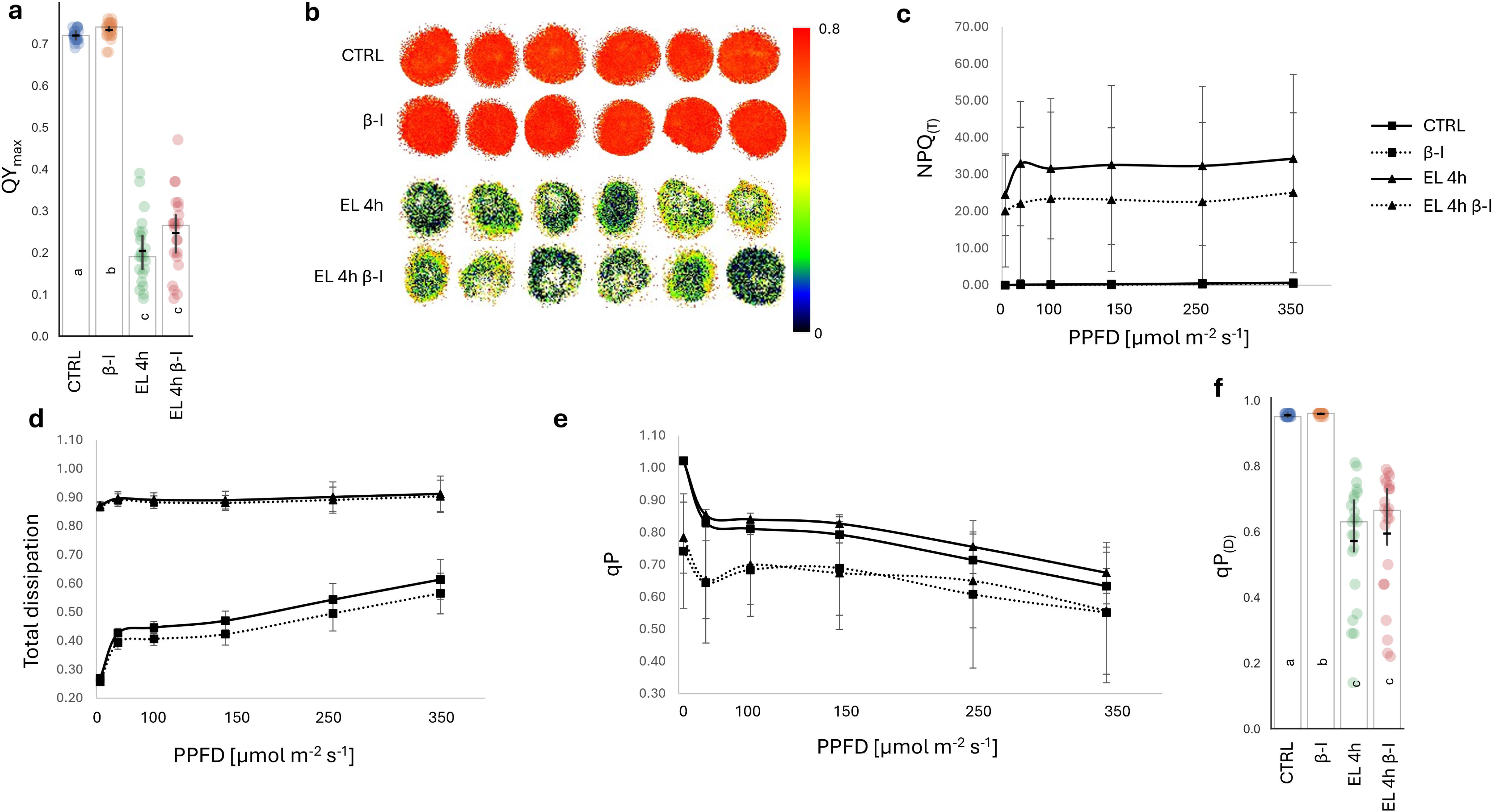
The detrimental photosynthetic response is specific to β-cyclocitral and is not reproduced by β-ionone. 28-day after inoculation Physcomitrium plants were pretreated for 4 h with volatile β-ionone (β-I; 50 µL) or control conditions and then exposed, where indicated, to EL for 4 h. (a) QY_max_ under control, β-I, EL and EL + β-I conditions. (b) Representative QY_max_ images. (c) NPQ_(T)_. (d) Total energy dissipation. (e) qP during the light-response curve. (f) qP_(D)_ after 1 min of darkness. Data are means ± SD. Asterisks indicate significant differences relative to the corresponding control. Alternatively, different letters are assigned to statistically different groups.

### 3.4 β-cc accumulates under EL in Physcomitrium

Recently, Rieseberg et al. performed a comprehensive study of stress response kinetics in three streptophyte species spanning approximately 600 million years of evolutionary divergence, including Physcomitrium (Rieseberg *et al*., 2025). They observed a rapid accumulation of β-cc in Physcomitrium under EL (800-1050 μmol photons m^−2^ s^−1^), reaching levels up to three times those observed under control conditions after 1 h of stress. Interestingly, β-cc accumulation under EL was not observed in the filamentous alga *Zygnema circumcarinatum*.

We measured the concentrations of β-cc, β-cyclogeranic acid (β-cca) and β-I under our stress conditions. 4 h of EL did not induce a detectable increase in apocarotenoid levels, whereas prolonged exposure did (16 h, Figure 4a). β-cc increased approximately sixfold from approximately 2 to 12 ng g^−1^ fresh weight (FW), whereas β-cca and β-I showed smaller increases (Figure 4a). Thus, apocarotenoid accumulation became detectable only after prolonged EL exposure, despite the marked inhibition of photosynthesis observed after 4 h.

**Figure 4:**
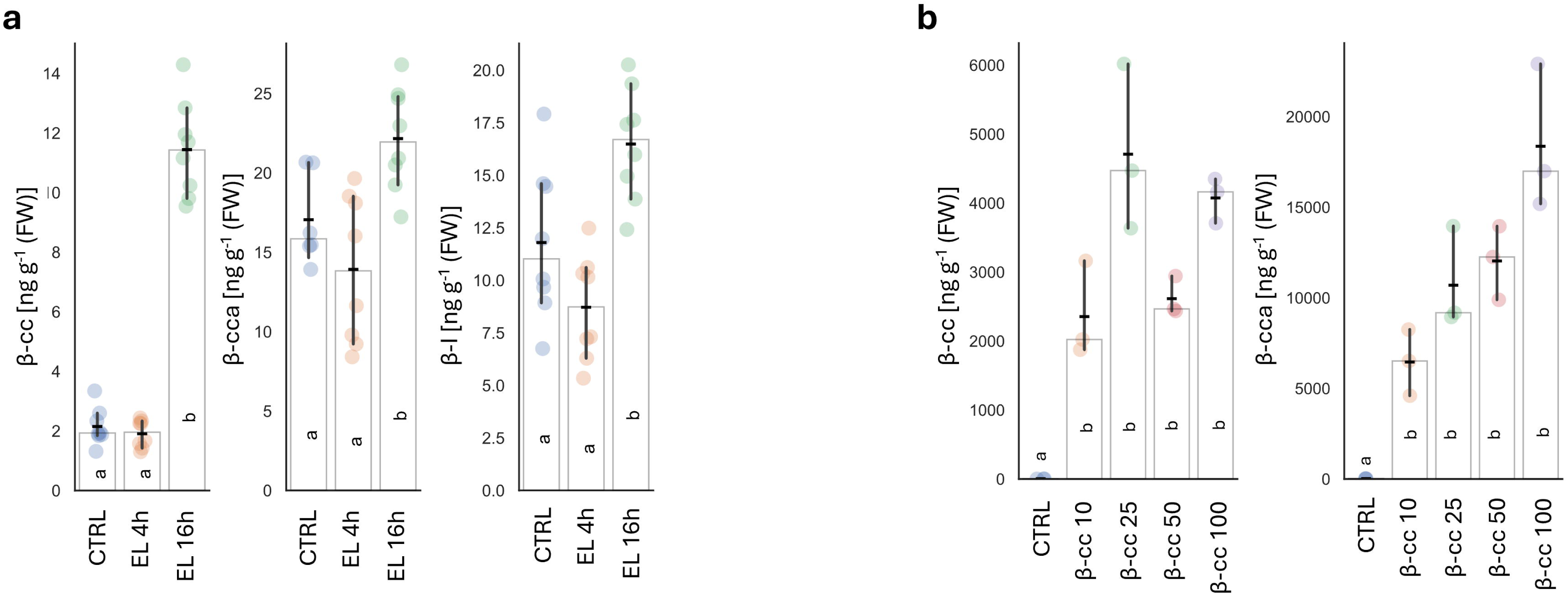
β-cyclocitral and β-cyclocitric acid accumulate in Physcomitrium under excess light and after volatile β-cyclocitral treatment. Endogenous and treatment-induced apocarotenoid levels were quantified by GC-MS. (a) Concentrations of β-cc, β-cyclocitric acid (β-cca) and β-ionone (β-I) in control samples and after 4 or 16 h of EL. (b) β-cc and β-cca accumulation after 4 h exposure to increasing volumes of volatile β-cc. Data are means ± CI95%. Different letters are assigned to statistically different groups.

We also quantified β-cc and its derivative, β-cca, in β-cc-treated samples and found that all treatments caused a marked increase in tissue β-cc and β-cca concentrations in Physcomitrium (Figure 4b). While β-cc levels appeared to plateau at approximately 2000–4000 ng g^−1^ FW across treatment volumes, β-cca showed a more linear response, with concentrations increasing from approximately 6000 to 18000 ng g^−1^ FW. These concentrations greatly exceeded endogenous levels measured under EL. Treatment-induced accumulation was also more pronounced than in Arabidopsis, where concentrations increased approximately 100-fold above physiological levels (Carna *et al*., 2026).

In light of these results, we tested whether a longer EL treatment or a lower volume of β-cc could modify our observations. After 16 h of EL, β-cc still provided no detectable protection in Physcomitrium (Supplemental Figure S2a, b, c). Regarding the lower treatment volume, we pretreated plants with 2 μL of β-cc. We chose not to dilute β-cc to avoid introducing solvent-dependent differences in volatilization within the closed desiccator. Even the 2-μL treatment induced a small but significant reduction in QY_max_ and affected NPQ induction, as observed at higher β-cc treatment volumes (Supplemental Figure S2d and e). This treatment likewise failed to provide detectable protection against EL in Physcomitrium (4 h, Supplemental Figure S2f and g).

**Figure S2:**
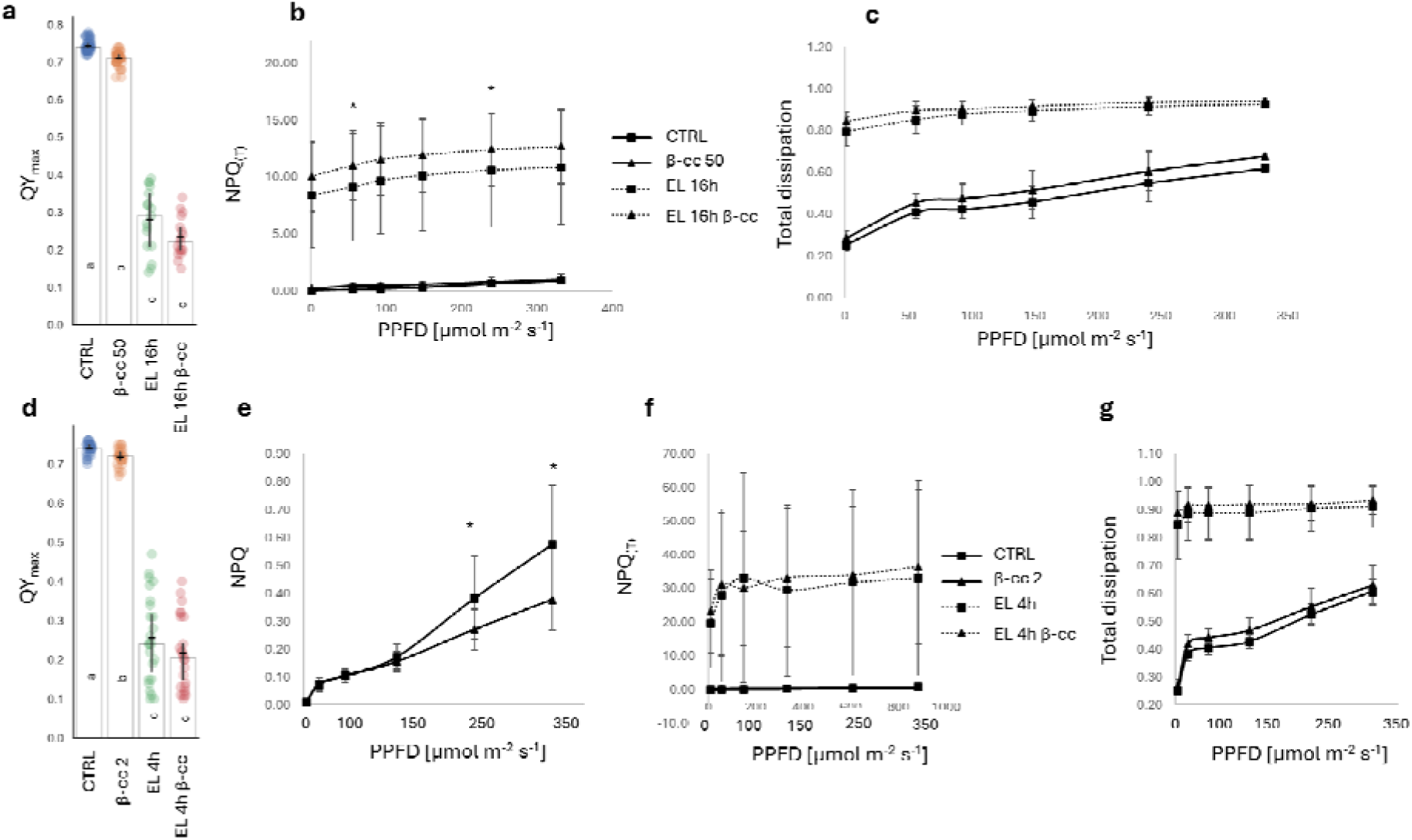
β-cyclocitral does not protect Physcomitrium from photoinhibition at lower treatment or longer stress conditions. 28-day after inoculation Physcomitrium plants were pretreated with β-cc and exposed to EL for 4 or 16 h. Panels show QY_max_, NPQ, NPQ_(T)_ and total energy dissipation in experiments with the low-volume β-cc treatment or 16 h EL. Data are means ± SD. Asterisks indicate significant differences relative to the corresponding control. Alternatively, different letters are assigned to statistically different groups.

### 3.5 β-cc does not alter the abundance of photosynthetic proteins

To investigate the basis of the detrimental effects of β-cc, we examined the abundance of selected proteins involved in photosynthesis. These included the PSII core protein D2 and the antenna proteins LHCII, the NPQ-related proteins PsbS and LHCSRs, both of which are present in Physcomitrium, and Rubisco. None of the proteins analyzed showed a significant difference in abundance between β-cc-treated and untreated samples (Figure 5a and b). However, the 4-h treatment may have been too short to produce detectable changes in protein abundance. The observed physiological effects could instead involve post-translational regulations.

**Figure 5:**
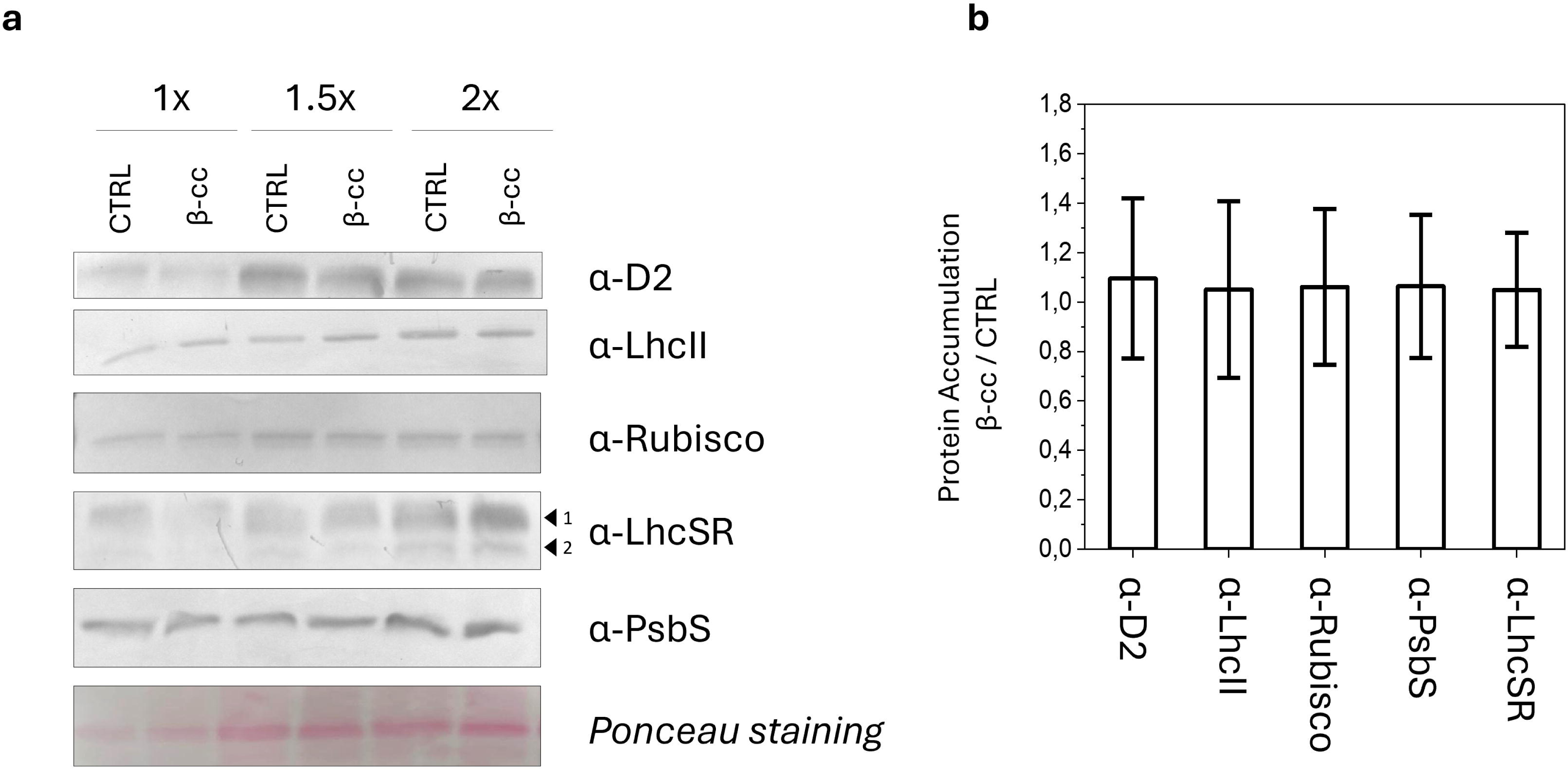
β-cyclocitral does not alter the abundance of photosynthetic proteins. Total protein extracts from Physcomitrium plants exposed for 4 h to β-cc (50 µL) or control conditions were analyzed by immunoblotting. (a) Representative immunoblots for D2, LHCII, Rubisco, LHCSR1 and 2 (black arrows) and PsbS; Ponceau staining is shown as loading control. (b) Densitometric quantification of protein accumulation in β-cc-treated samples relative to controls. Data are means ± SD. No significant differences were detected under the conditions tested.

### 3.6 β-cc increases lipid peroxidation in Physcomitrium

Impaired photosynthetic performance can be associated with oxidative damage. To assess whether the effects of β-cc were accompanied by increased lipid peroxidation, we quantified malondialdehyde (MDA) as a proxy for lipid peroxidation. In contrast to the response reported in Arabidopsis, β-cc treatment induced a small but significant increase in MDA content even under control conditions (D’Alessandro *et al*., 2018). This increase was more pronounced after 4 and 16 h of EL, when β-cc-treated samples contained approximately twice as much MDA as untreated samples (Figure 6a).

**Figure 6:**
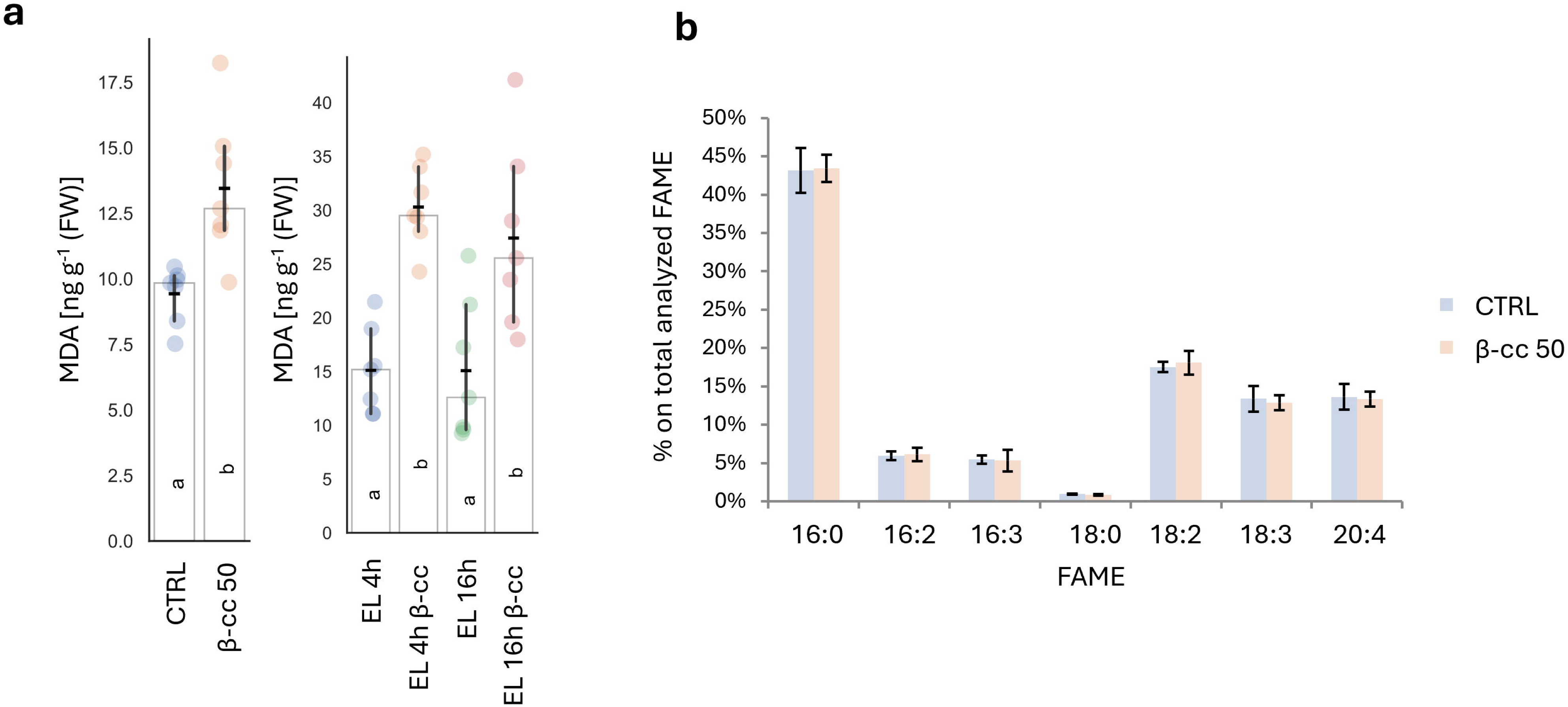
β-cyclocitral increases lipid peroxidation without altering the relative fatty-acid profile. 28-day after inoculation Physcomitrium plants were exposed to β-cc (50 µL) or control conditions and, where indicated, to EL for 4 or 16 h. (a) Malondialdehyde (MDA) content, used as a proxy for lipid peroxidation. (b) Relative abundance of identified fatty acid methyl esters (FAMEs) in control and β-cc-treated samples. Values in panel b are expressed as percentage of total identified FAMEs. Data are means ± SD. Different letters are assigned to statistically different groups.

To investigate the origin of β-cc-induced oxidation, we analyzed fatty acid methyl esters (FAMEs). Because fatty acids are differentially distributed among organellar membranes, changes in their profiles could provide clues to the cellular membranes affected (Haslam *et al*., 2024). However, we detected no differences in the relative abundance of the identified fatty acids (16:0, 16:2, 16:3, 18:0, 18:2, 18:3 and 20:4) between treated and untreated samples (Figure 6b).

### 3.7 Transcriptional responses to β-cc and EL

β-cc reduced photosynthetic performance in Physcomitrium after only 4 h of exposure, even at a treatment volume of 2 μL. β-cc treatment was also associated with increased lipid peroxidation. As the biochemical analyses did not fully explain why β-cc failed to confer protection against EL in Physcomitrium, we performed RNA-seq analysis to characterize the transcriptional responses to β-cc and EL.

β-cc elicited a strong transcriptional response comprising 2102 upregulated genes and 618 downregulated genes relative to the control, demonstrating extensive transcriptional reprogramming in response to β-cc treatment (Figure 7a). EL exposure (16 h) also induced a substantial transcriptional response, with 2693 upregulated genes and 3345 downregulated genes, broadly consistent with the extensive response reported previously (around 2500 upregulated genes and 2000 downregulated genes after 6 h of EL) (Rieseberg *et al*., 2025). Principal component analysis (PCA) of the 6000 most variable genes, a number selected based on the number of differentially expressed genes (DEGs), revealed a clear separation of the β-cc-treated and EL-exposed samples from control samples, as well as a distinction between the two treatments (Figure 7b). The overlap between the responses to β-cc and EL was also evident in the Venn diagram (Figure 7c), which showed that approximately 66% of genes induced in response to β-cc were also induced under EL although EL downregulated a substantially larger number of genes. The similarity between the two transcriptional responses was further supported by heatmaps of the most variable genes and sample-to-sample distances, which grouped β-cc-treated samples with EL-exposed samples (Supplemental Data 1).

**Figure 7:**
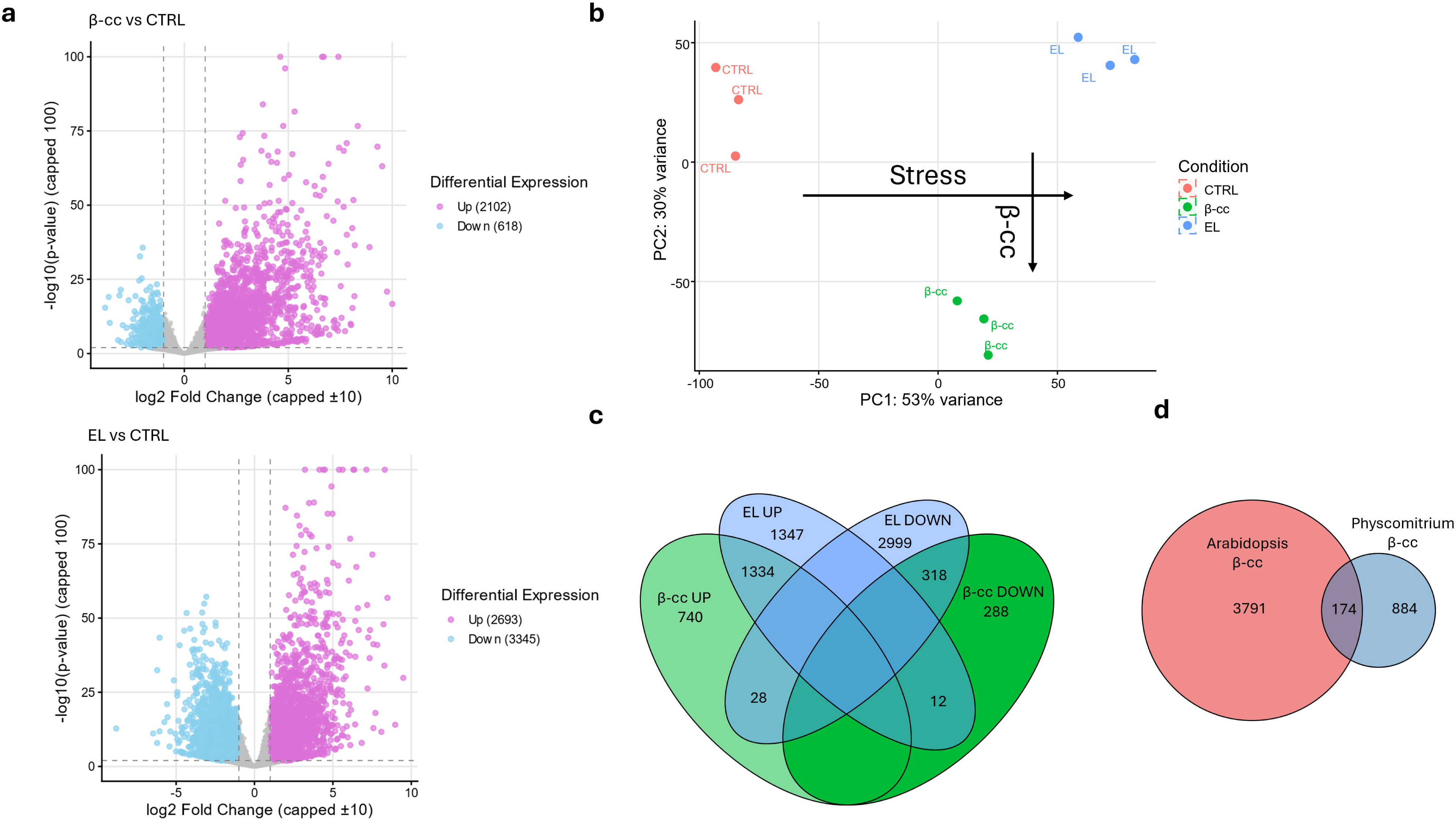
β-cyclocitral activates a broad stress-associated transcriptional response that overlaps with the excess-light response. RNA-seq analysis was performed on Physcomitrium plants treated with β-cc or exposed to EL. (a) Volcano plots showing differentially expressed genes in β-cc-treated samples and in EL-exposed samples relative to controls. (b) Principal component analysis of the 6000 most variable genes. (c) Overlap among upregulated and downregulated genes in response to β-cc and EL. (d) Overlap between β-cc-responsive genes in Physcomitrium and Arabidopsis after mapping to Arabidopsis homologs (multiblast). Differentially expressed genes were defined using the thresholds reported in Materials and Methods (p-adj < 0.01 and |Log2|>1).

Gene Ontology, KEGG, and MapMan analyses (DEG-based and GSEA analyses) were performed using both functional annotation for Physcomitrium (Bi *et al*., 2024) and annotations for homologous genes in Arabidopsis (Ashburner *et al*., 2000; Thimm *et al*., 2004; Subramanian *et al*., 2005; Kanehisa *et al*., 2010). Among the selected GO terms, “response to heat” showed the strongest enrichment in genes upregulated by β-cc, consistent with its enrichment under EL. Other enriched terms included responses to high light intensity, hydrogen peroxide and water deprivation. MapMan analysis also identified enrichment of DREB-type transcription factors among upregulated genes, whereas GSEA showed negative enrichment of the cell cycle gene set (Table 1, Supplemental Data 1). Together, these enrichments indicate that β-cc treatment activates a broad stress-associated transcriptional response in Physcomitrium. In addition, we observed enrichment of the ABC1 family of atypical kinases, involved in tocopherol synthesis and stress response (Lundquist *et al*., 2012), among upregulated genes and of GRAS transcription factors among downregulated genes. The latter observation is noteworthy because SCL14, a member of the GRAS family, participates in β-cc signaling in Arabidopsis (Table 1, Supplemental Data 1).

**Table 1:** selected functional enrichments induced by β-cc.

| GO code | GO term | Score | Gene count | Algorithm |
| --- | --- | --- | --- | --- |
| GO:0009408 | response to heat | 12.7 | 87 | enrichGO |
| GO:0009644 | response to high light intensity | 9.91 | 54 | enrichGO |
| GO:0042542 | response to hydrogen peroxide | 7.58 | 33 | enrichGO |
| GO:0009414 | response to water deprivation | 5.4 | 68 | enrichGO |
| GO:2001141 | regulation of RNA biosynthetic process | -8.51 | 185 | enrichGO |
| GO:0006468 | protein phosphorylation | -6.22 | 139 | enrichGO |
| GO:0009903 | chloroplast avoidance movement | -5 | 10 | enrichGO |
| 15.7.7.2 | DREB-type transcription factor | 4.2 | 20 | Enricher:MapMan |
| 18.8.7.4 | ABC1 kinase | 4 | 10 | Enricher:MapMan |
| 15.7.12 | GRAS transcription factor | -4 | 9 | Enricher:MapMan |
| GO:0007049 | cell cycle | -1.67 |  | GSEA |

EL-exposed samples also showed enrichment of stress-related GO terms, including responses to chemicals, heat and abscisic acid (Table 2). The main differences between the β-cc and EL responses concerned photosynthesis: genes downregulated under EL were enriched for GO terms related to photosynthesis, light harvesting and chlorophyll biosynthesis. A similar difference between β-cc and EL in the transcriptional regulation of photosynthesis has also been observed in Arabidopsis (Carna *et al*., 2026).

**Table 2:** selected functional enrichments induced by EL.

| GO code | GO term | Score | Gene count | Algorithm |
| --- | --- | --- | --- | --- |
| GO:0042221 | response to chemical | 8.3 | 628 | enrichGO |
| GO:0009408 | response to heat | 7.2 | 89 | enrichGO |
| GO:0009737 | response to abscisic acid | 6.9 | 158 | enrichGO |
| GO:0015979 | photosynthesis | -17.6 | 161 | enrichGO |
| GO:0009765 | light harvesting | -11.6 | 56 | enrichGO |
| GO:0015995 | chlorophyll biosynthetic process | -8.3 | 24 | enrichGO |
| GO:0007049 | cell cycle | -2.2 |  | GSEA |

We compared the β-cc transcriptional response in *Physcomitrium* with that reported in *Arabidopsis* (Carna *et al*., 2026). Among 1,058 *Physcomitrium* β-cc-responsive genes with assigned *Arabidopsis* homologues, 174 (16.4%) were shared between the two responses (Figure 7d). To avoid dependence on homologue assignment, we compared β-cc- and EL-induced responses within each species using rank-overlap enrichment (Figure S3). *Arabidopsis* showed limited concordance among strongly upregulated genes and pronounced discordance across broader rank fractions, whereas *Physcomitrium* displayed strong concordance throughout the analyzed rankings.

**Figure S3:**
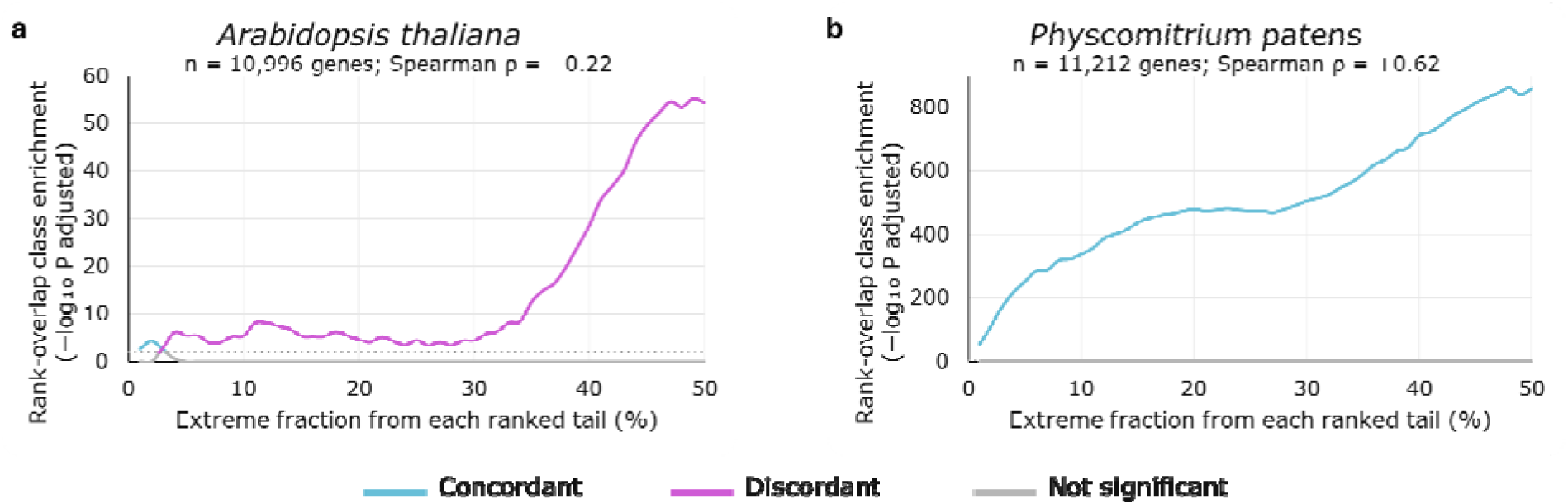
Rank-overlap enrichment of β-cyclocitral- and excess-light-induced transcriptional responses in Arabidopsis and *Physcomitrium*. Concordant and discordant transcriptional responses to β-cc and excess light in (a) *A. thaliana* and (b) *P. patens*. Enrichment is expressed as −log_10_(adjusted P) across increasing fractions of the ranked gene lists.

### 3.8 Limited conservation of the Arabidopsis β-cc-associated detoxification in Physcomitrium

Two major mechanisms have been implicated in β-cc-induced tolerance to EL in *Arabidopsis thaliana*: one involving METHYLENE BLUE SENSITIVITY 1 (MBS1) and the other involving the GRAS protein SCARECROW-LIKE 14 (SCL14) (Shumbe *et al*., 2017; D’Alessandro *et al*., 2018). MBS1 contributes to singlet-oxygen (^1^O_2_)-responsive gene regulation and has recently also been shown to undergo a ^1^O_2_-dependent phase transition, forming chloroplast-associated condensates that attenuate light penetration and protect chloroplasts from photodamage (Shumbe *et al*., 2017; Shao *et al*., 2026). SCL14 regulates the TGAII-dependent induction of the transcription factor ANAC102, which controls three other ANAC transcription factors (ANAC002, ANAC032 and ANAC081). This regulatory cascade induces the expression of numerous enzymes involved in xenobiotic detoxification (Fode *et al*., 2008; D’Alessandro *et al*., 2018; Cresta & D’Alessandro, 2023).

We initially examined the conservation of these components in Physcomitrium using the available multiblast-based homology assignments and integrated these results with the transcriptional responses to β-cc and EL (Goodstein *et al*., 2012; Bi *et al*., 2024). As shown in Table 3, this analysis identified three putative homologs of MBS1 and three of SCL14. However, no putative homologs were assigned to ANAC102, a key regulator of the downstream transcriptional cascade, or to ANAC032 and ANAC081. ANAC002 was the only member of this four-factor ANAC module with putative homologs identified in Physcomitrium. Whereas ANAC002 was strongly induced by β-cc in Arabidopsis, its putative Physcomitrium homologs were induced by EL but not by β-cc (Table 3).

**Table 3:** putative homologs of the detoxification pathway in Physcomitrium based on multiblast comparisons. Numbers are log_2_ fold changes (Ns = not significantly differentially expressed).

| Name | TAIR ID | *Physcomitrium* | Pp β-cc | At β-cc | Pp EL |
| --- | --- | --- | --- | --- | --- |
| MBS1 | AT3G02790 | PpV7_09g0004900 | Ns | Ns | Ns |
|  |  | PpV7_15g0005700 | Ns |  | Ns |
|  |  | PpV7_26g0007300 | -0.9 |  | Ns |
| SCL14 | AT1G07530 | PpV7_17g0059800 | Ns | Ns | 2.4 |
|  |  | PpV7_02g0060700 | Ns |  | Ns |
|  |  | PpV7_19g0076400 | Ns |  | 1.1 |
| ANAC102 | AT5G63790 | No homologs |  | 3.3 |  |
| ANAC002 | AT1G01720 | PpV7_03g0039900 | Ns | 3.9 | 3.6 |
|  |  | PpV7_13g0044700 | Ns |  | 1.5 |
| ANAC032 | AT1G77450 | No homologs |  | 2.5 |  |
| ANAC081 | AT5G08790 | No homologs |  | 2.4 |  |
| CYP81D11 | AT3G28740 | No homologs |  | 6.4 |  |
| AER | AT5G16970 | No homologs |  | 2.6 |  |
| AKR4C | AT2G37760 | No homologs |  | C8 1.1 |  |
|  | AT2G37770 | PpV7_10g0065900 | 6 | C9 Ns | 5 |
|  |  | PpV7_12g0019300 | Ns |  | Ns |
| Chl-ADR | AT3G04000 | No homologs |  | 2 |  |
|  | AT1G54870 | PpV7_26g0065400 | 4.9 | Ns | 4.2 |
|  |  | PpV7_18g0049800 | 4.8 |  | 4.1 |
| SDR1 | AT3G61220 | No homologs |  | 1.5 |  |
| GST | AT2G29490 | No homologs |  | U1 5 |  |
|  | AT2G29470 | No homologs |  | U3 2.1 |  |
|  | AT2G29420 | No homologs |  | U7 5.1 |  |
|  | AT3G09270 | No homologs |  | U8 2.6 |  |
|  | AT1G78380 | No homologs |  | U19 2.6 |  |
|  | AT1G17170 | No homologs |  | U24 4.5 |  |
|  | AT2G02390 | PpV7_01g0158300 | Ns | Z1 2 | Ns |

The same homology-based analysis identified putative counterparts of only a few Arabidopsis marker genes for SCL14-dependent xenobiotic detoxification in Physcomitrium: two aldo-keto reductase (AKR) 4C genes, one of which was induced by β-cc and EL, two chloroplast aldehyde reductases (Chl-ADR) genes, both induced by β-cc and EL, and one glutathione S-transferase (GST) gene, which was not induced by either treatment (Table 3).

To distinguish broad protein-family conservation from conservation of the specific β-cc-associated subgroups, we subsequently performed a comparative phylogenomic analysis across 14 photosynthetic organisms, selected to represent major photosynthetic lineages and to include species with documented β-cc production.

This analysis revealed markedly different evolutionary patterns among the components of the pathway. MBS1 represented the clearest case of deep conservation: three Physcomitrium proteins were recovered within the compact MBS1 orthogroup and shared approximately 70–73% amino-acid identity with *A. thaliana* MBS1 (Table 4, Supplemental Figure S4). Interestingly, the Physcomitrium MBS1-like proteins grouped with homologs from other non-seed-plant lineages, including streptophyte algae, rather than with angiosperm homologs.

**Table 4:**
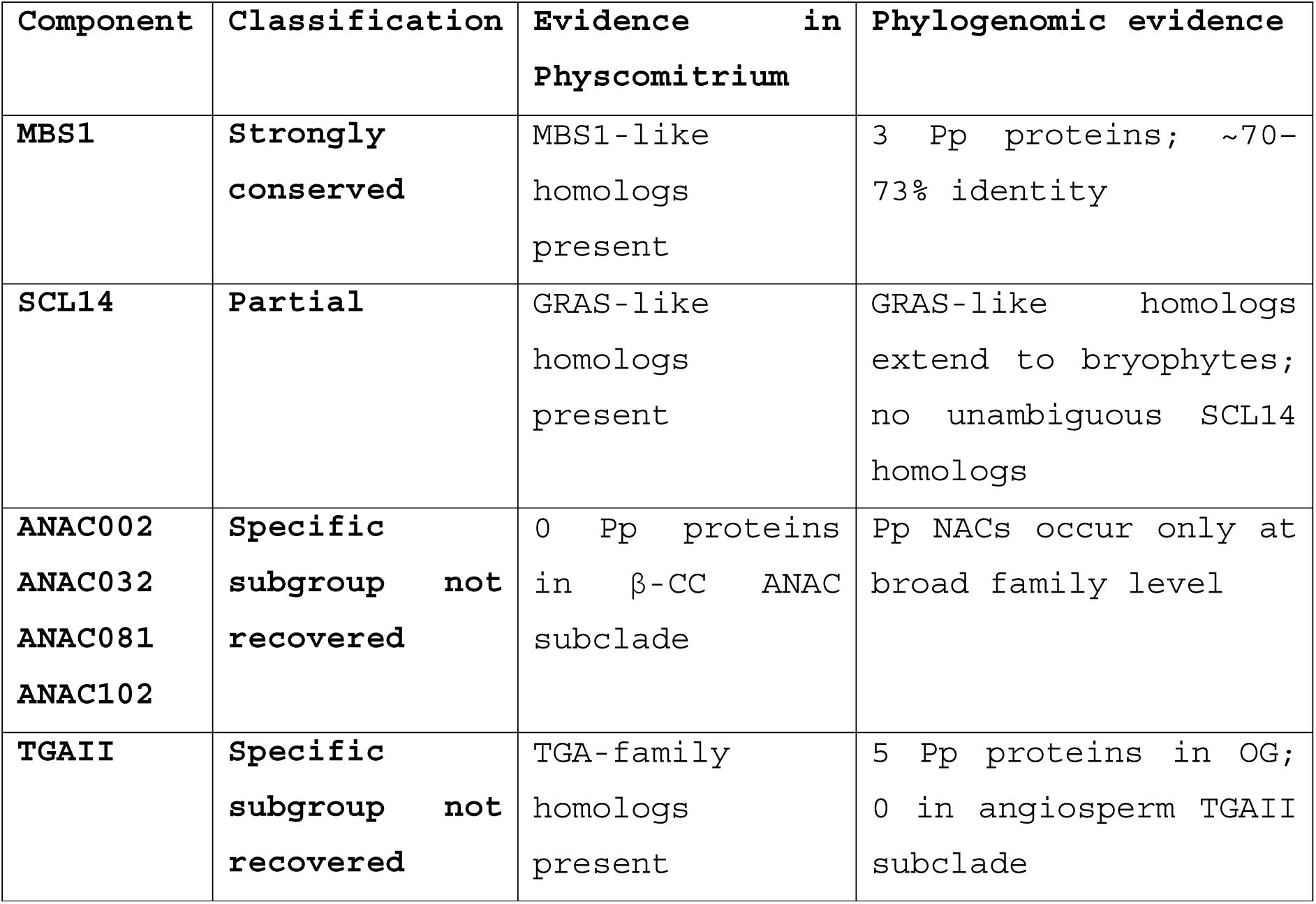
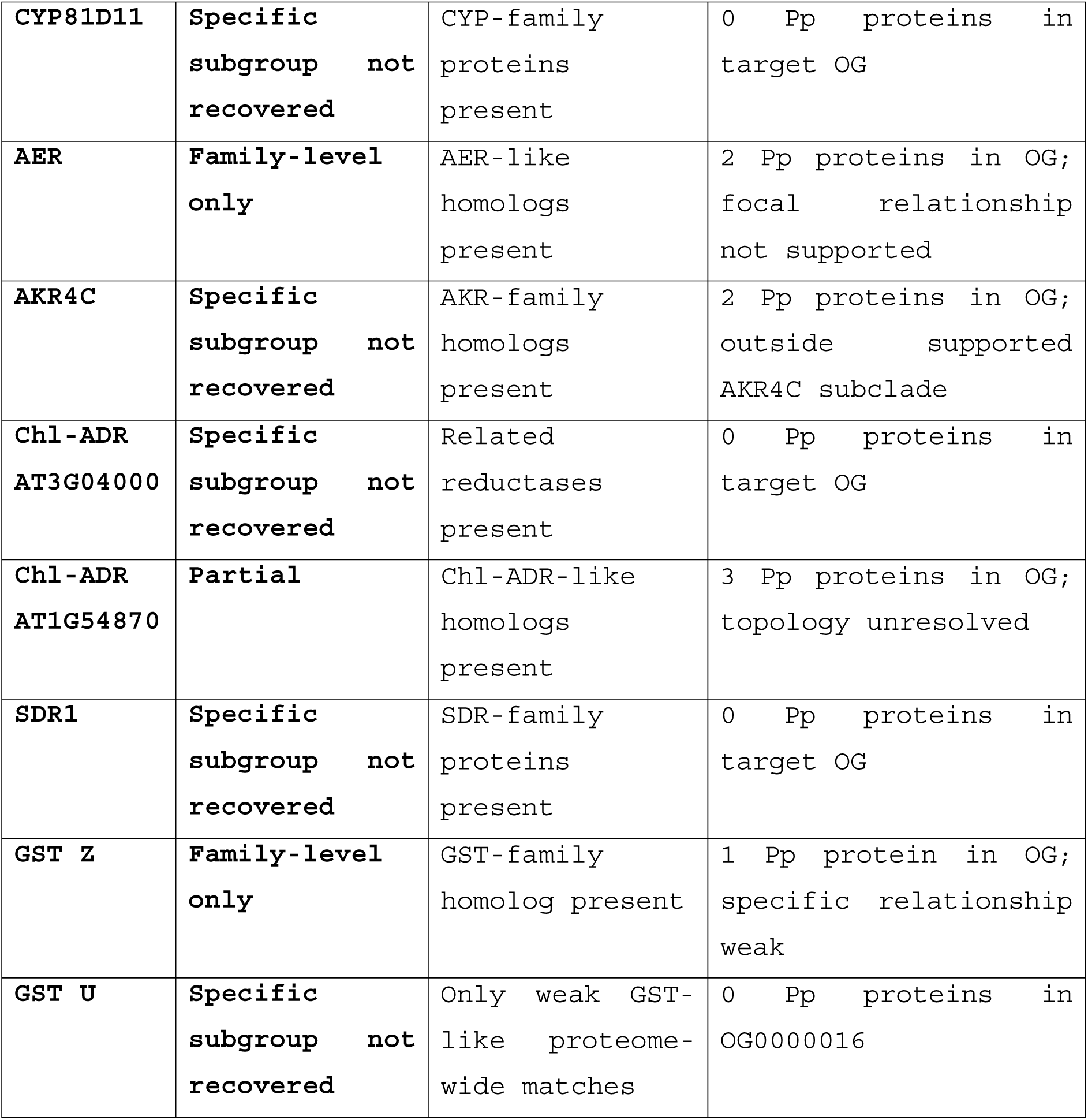
putative homologs of the detoxification pathway in Physcomitrium based on phylogenetic evidence.

**Figure S4:**
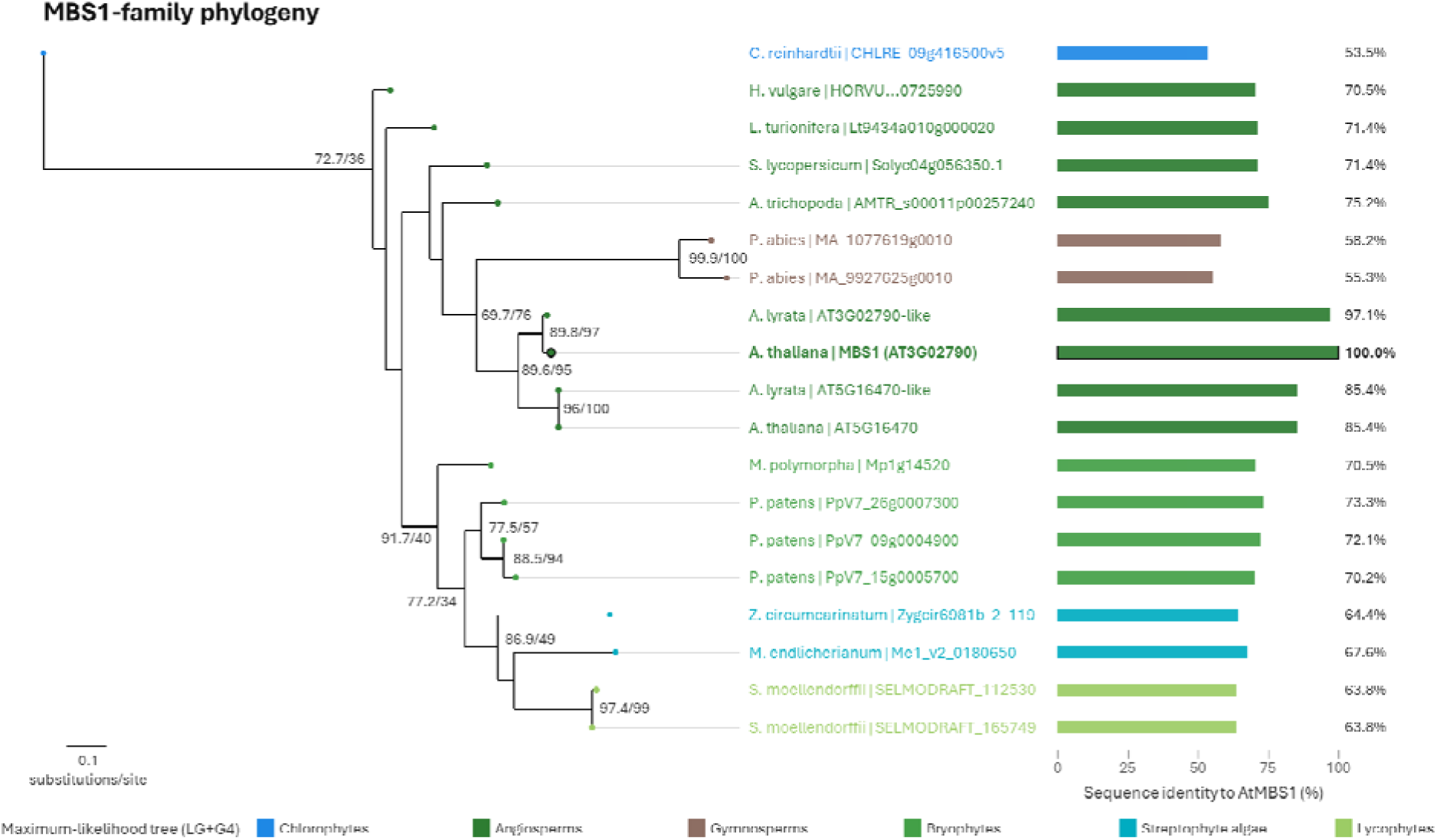
MBS1-family phylogeny across photosynthetic lineages. Maximum-likelihood phylogeny of the MBS1-containing orthogroup inferred from representative proteomes. Branch labels indicate SH-aLRT/UFBoot support where shown. Bar plots indicate amino-acid sequence identity to Arabidopsis MBS1. Taxa are coloured by major lineage.

By contrast, the SCL14-containing family showed greater sequence divergence, already among Arabidopsis paralogs, and the phylogenetic analysis did not resolve an unambiguous relationship between the Physcomitrium proteins and the specific lineage containing Arabidopsis SCL14 (Supplemental Figure S5).

**Figure S5:**
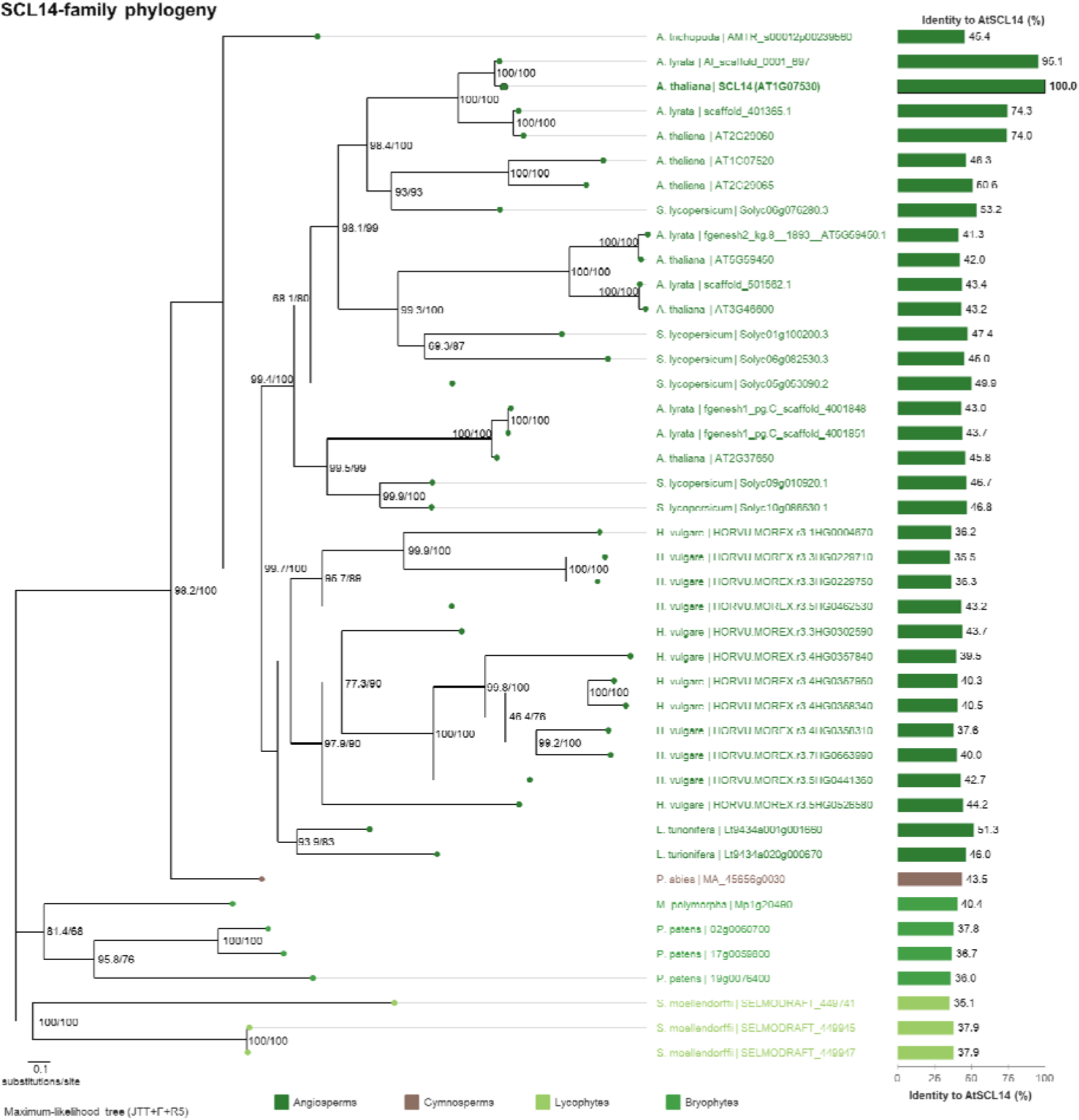
SCL14-family phylogeny across photosynthetic lineages. Maximum-likelihood phylogeny of the SCL14-containing GRAS-family orthogroup. Branch labels indicate SH-aLRT/UFBoot support where shown. Bar plots indicate amino-acid sequence identity to Arabidopsis SCL14. Physcomitrium proteins are recovered within the broader family but not unambiguously within the Arabidopsis SCL14 target lineage.

The strongest divergence was observed in the ANAC regulatory module. ANAC081 and ANAC102 formed a well-supported β-cc-associated NAC subclade together with ANAC002 and ANAC032 (ATAF subfamily) (Takasaki *et al*., 2015), but no Physcomitrium protein was recovered within this group. Physcomitrium NAC proteins were detected only at the broader NAC-family level, indicating conservation of the NAC family itself without recovery of members of the specific regulatory subgroup associated with β-cc-induced detoxification in Arabidopsis (Table 4, Supplemental Figure S6). Within angiosperms, the ANAC081/ANAC102-related branch was also not recovered in *L. turionifera*, although this species retained candidates associated with the ANAC002 branches. By contrast, *H. vulgare* proteins were recovered within the ANAC081/ANAC102-related lineage (Supplemental Figure S6, S7). We further focused on this evolutionary node by adding more monocotyledon species to the analyses, and confirmed that while terrestrial species such as wheat, rice and *Brachypodium distachyon*, show an element of the ANAC081/ANAC102-related lineage, aquatic species such as *Zostera marina* and *Spirodela polyrhiza* developed substantial evolutionary divergence from this clade (Supplemental Figure S8). Thus, the distribution of ANAC081/ANAC102-like proteins differed between the terrestrial and aquatic monocot species sampled here, whereas Physcomitrium lacked identifiable members of the broader β-cc-associated ATAF subgroup.

**Figure S6:**
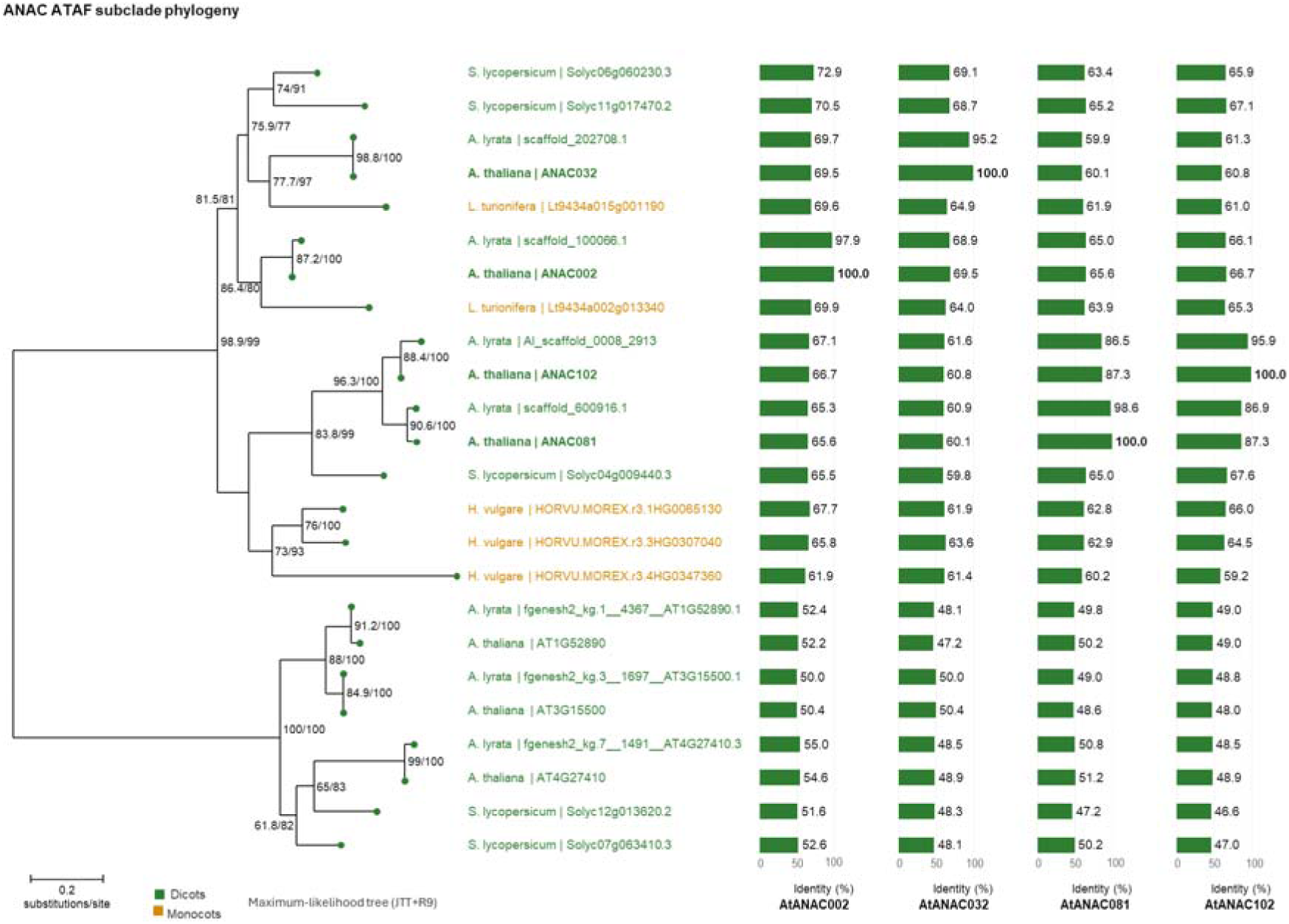
Phylogeny of the ANAC ATAF subclade associated with β-cyclocitral responses. Maximum-likelihood phylogeny of ANAC002/ANAC032/ANAC081/ANAC102-related proteins. Branch labels indicate SH-aLRT/UFBoot support where shown. Bar plots show sequence identity to Arabidopsis ANAC002, ANAC032, ANAC081 and ANAC102. No Physcomitrium protein was recovered in the displayed β-cc-associated ATAF subclade.

**Figure S7:**
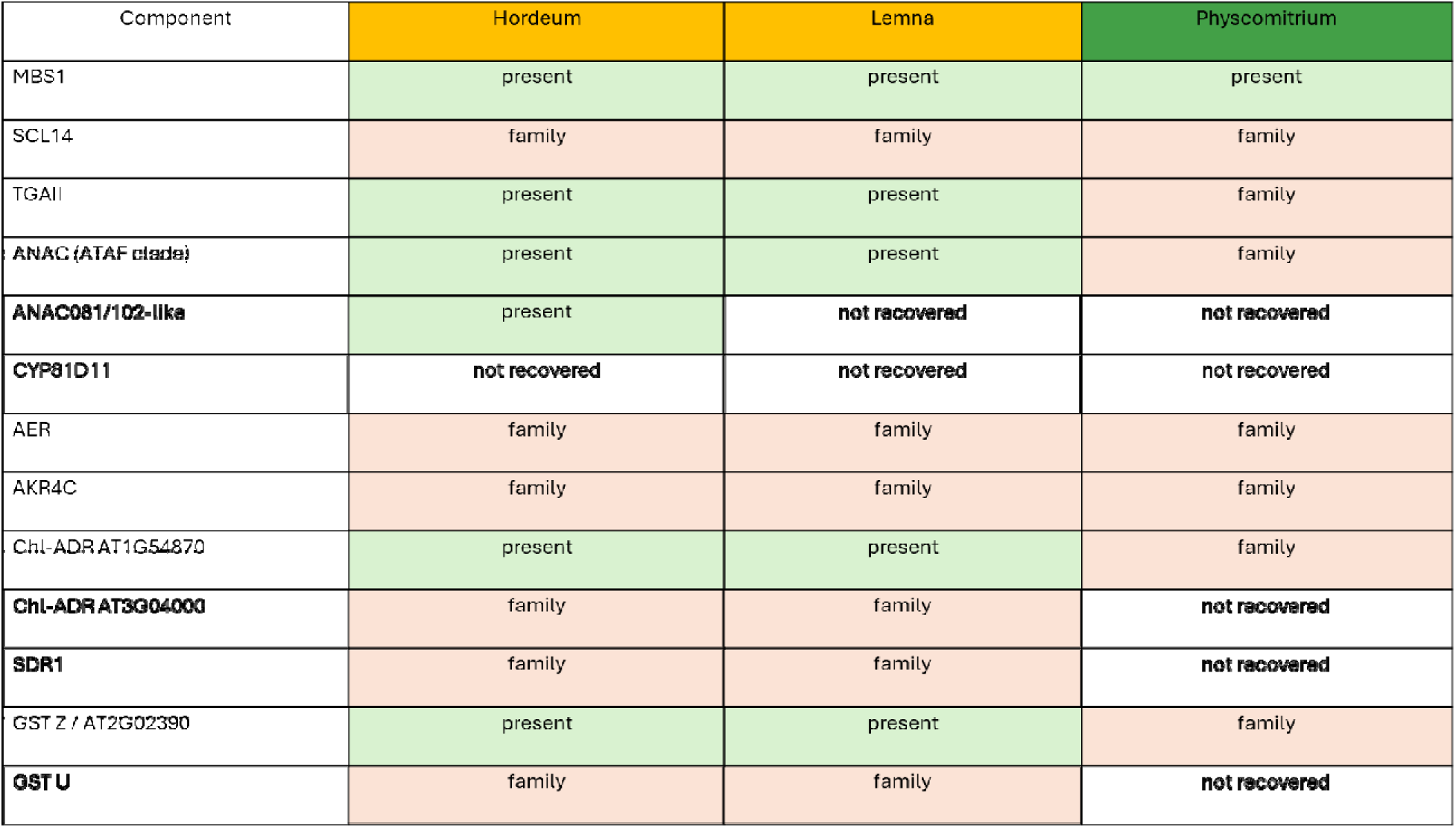
Summary of conservation of β-cyclocitral-associated regulatory and detoxification components. Classification of candidate components in Hordeum, Lemna and Physcomitrium. “Present” indicates that a protein from the species falls within the phylogenetic lineage containing the Arabidopsis reference group. “Family” indicates proteins in the same family but outside the specific Arabidopsis target lineage. “Not recovered” indicates that no member of the target orthogroup was recovered.

**Figure S8:**
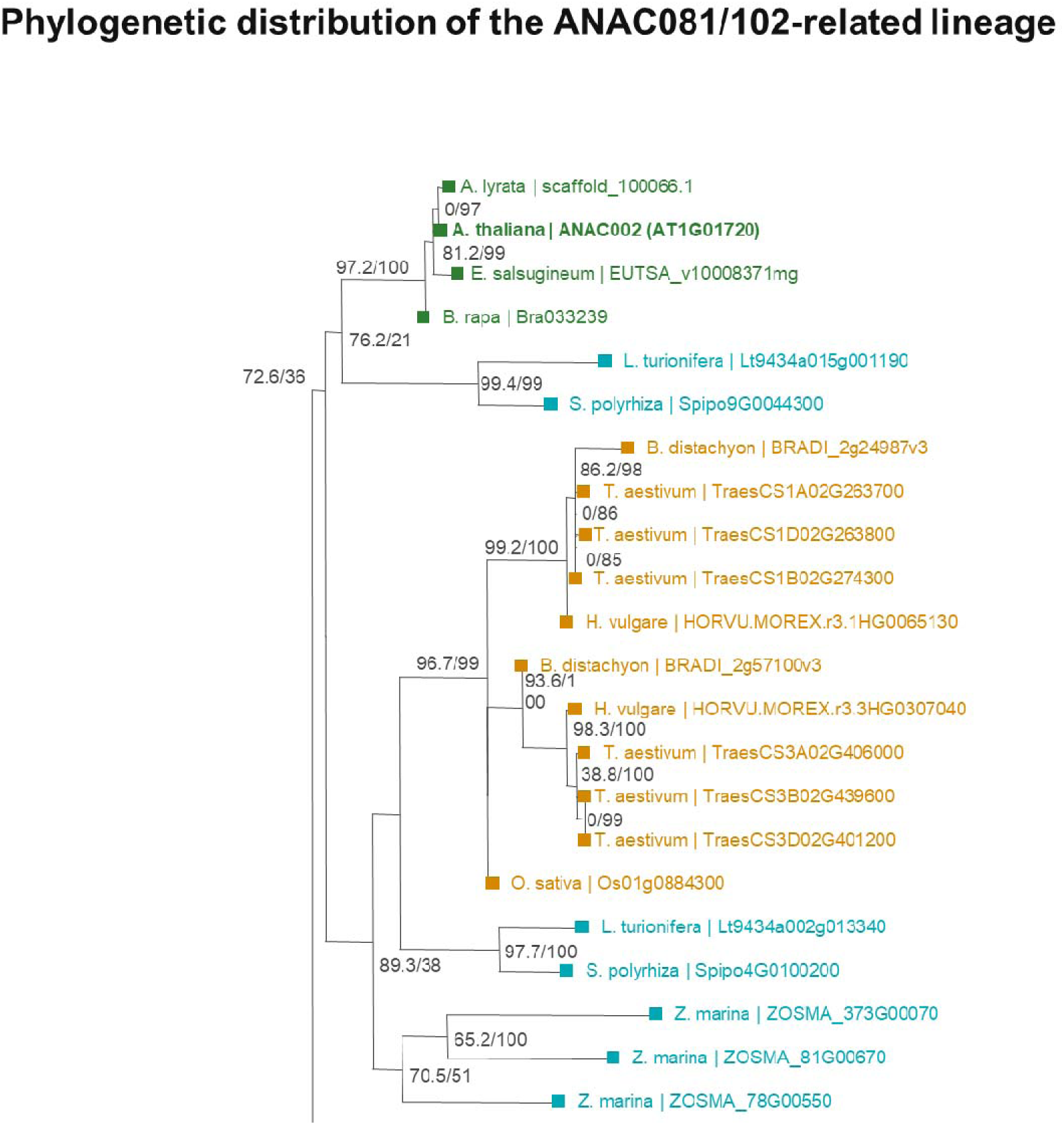
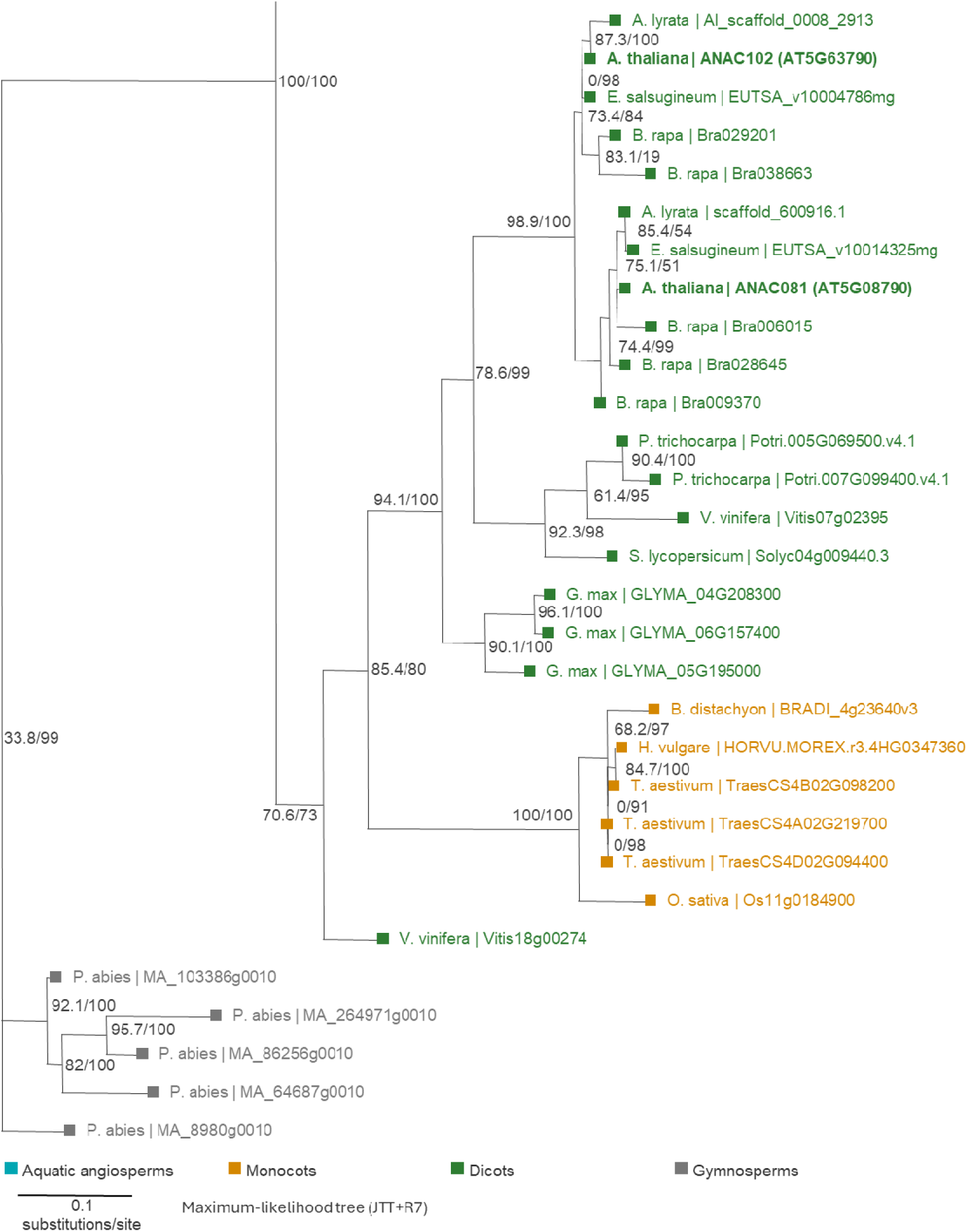
Phylogenetic distribution of the ANAC081/ANAC102-related lineage in angiosperms and gymnosperms. Maximum-likelihood phylogeny of the ANAC081/ANAC102-related lineage based on the NAC-core alignment after removal of ANAC032-related sequences for display clarity. Branch labels indicate SH-aLRT/UFBoot support where shown. Taxa are coloured by ecological group or major lineage as indicated.

A similar pattern was observed for several detoxification enzymes. Physcomitrium proteins belonging to the same orthogroups as AER, AKR4C and one Chl-ADR were identified, but phylogenetic analyses generally placed them outside the specific Arabidopsis target subclades (Figure S7). For CYP81D11, SDR1 and the GSTU subgroup, no Physcomitrium sequence was present in the corresponding target orthogroups. Proteome-wide similarity searches nevertheless detected related CYP, SDR, reductase and GST proteins in other orthogroups, demonstrating that the corresponding enzyme families are present in Physcomitrium, although members of the specific β-cc-associated subgroups were not recovered.

Together, these analyses indicate that the regulatory and enzymatic components of the Arabidopsis SCL14-dependent β-cc response are incompletely conserved in Physcomitrium and this divergence may contribute to the absence of detectable β-cc-induced protection against EL.

### 3.9 *Cis*-regulatory architecture of the β-cyclocitral response differs between Arabidopsis and Physcomitrium

In Arabidopsis, SCL14, together with class II TGA factors and ANAC transcription factors, regulates the transcriptional response to β-cc. Our phylogenetic analysis showed that SCL14 forms a distinct group from other LISCL proteins within *Brassicaceae*, as also observed for ANAC102 and ANAC081. Because these proteins regulate gene expression through promoter-associated mechanisms, we compared the *cis*-regulatory architecture of β-cc-induced genes in Arabidopsis and Physcomitrium. We first performed an unbiased survey of JASPAR plant transcription factor binding motifs across all β-cc-upregulated genes. Both species showed enrichment of multiple stress-associated *cis*-regulatory families, but the overall composition differed markedly (Figure 8a). In Arabidopsis, the global β-cc-responsive promoter set was enriched for several motif families, including NAC, bHLH/MYC/PIF, bZIP/ABRE, and TGA signatures. By contrast, the Physcomitrium response showed particularly strong enrichment of bZIP/ABRE and bHLH/MYC/PIF-related motifs, together with CAMTA, FAR1/FHY3, GRF, ARF, AP2/ERF, LBD, TCP and HSF-related signatures. We next asked which *cis*-regulatory signatures were preferentially associated with genes showing stronger induction (Figure 8b). In Arabidopsis, stronger responders (log2FC ≥ 1.5) were enriched for AP2/ERF, TGA, ARF/Auxin, bZIP/ABRE, TCP, TALE, bHLH/MYC and RVE/CCA1/LHY-related motifs, whereas they were depleted of MYB and Trihelix motifs. In Physcomitrium, only AP2/ERF-DREB and bZIP/ABRE families met the same enrichment criteria. These results indicate that TGA-associated signatures preferentially characterize the stronger Arabidopsis response.

**Figure 8:**
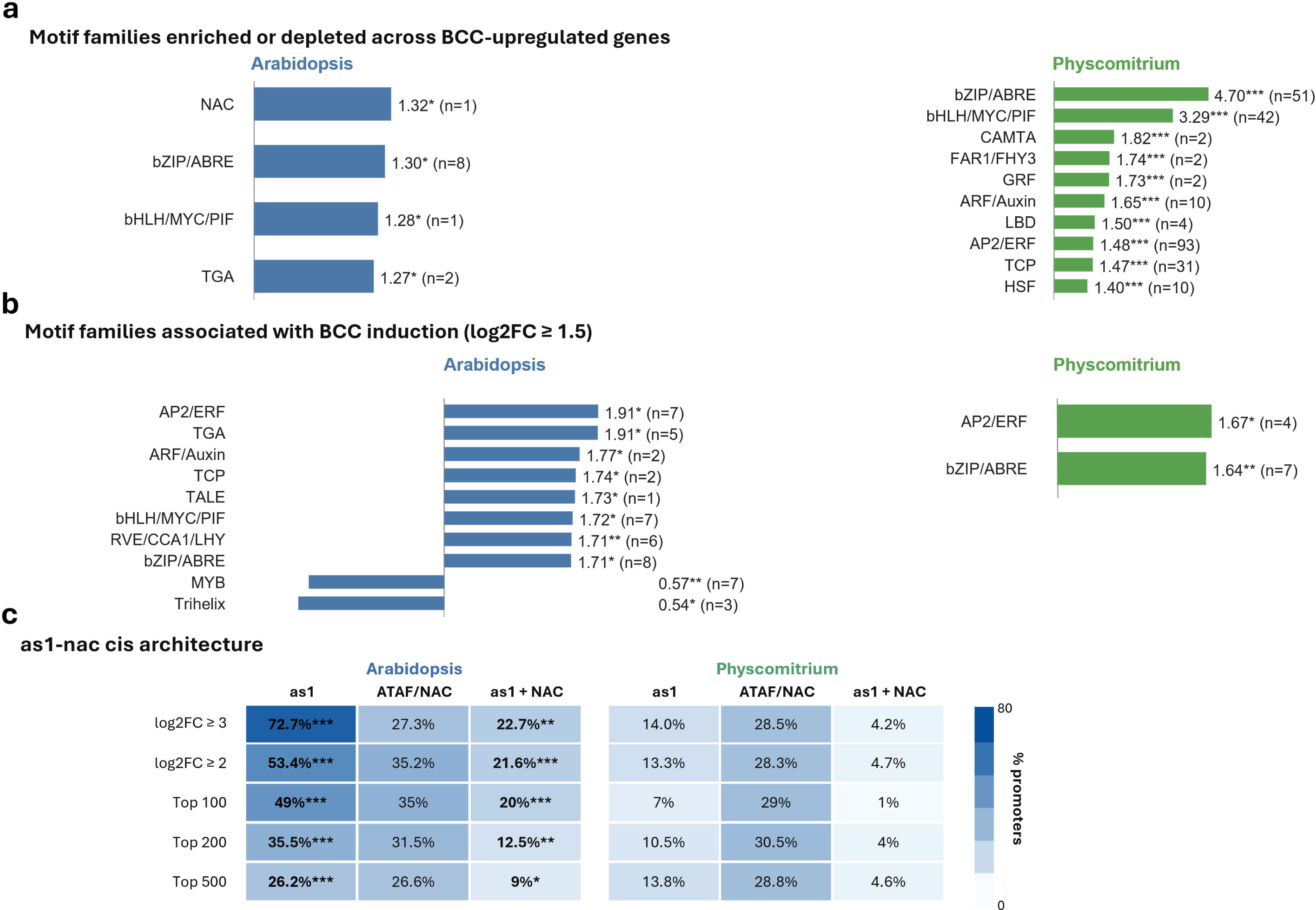
*Cis*-regulatory architecture of the β-cyclocitral response in Arabidopsis and Physcomitrium.=. (a) Enrichment of transcription factor motif families in promoters of β-cc-upregulated genes relative to background genes in *Arabidopsis thaliana* and *Physcomitrium patens*. Bars indicate the maximum odds ratio (OR) among significantly enriched JASPAR motifs assigned to each transcription factor family; numbers indicate the number of significant motifs within each family. Motifs with FDR < 0.05 and OR > 1 were considered enriched. (b) Motif families associated with stronger β-cc induction. Genes with log2FC ≥ 1.5 were compared with the remaining β-cc-upregulated genes within each species. Motif families shown contain at least one motif with FDR < 0.05 and OR ≥ 1.5. (c) Frequency of as-1-like, NAC-like and combined as1+NAC promoter architectures among genes showing increasing β-cc response amplitude. Heatmap cells indicate the percentage of promoters carrying each *cis*-regulatory feature among the top 500, 200 and 100 β-cc-induced genes ranked by log_2_FC and among genes with log_2_FC ≥ 2 or ≥ 3. ATAF/NAC represents the union of ANAC102-like, ANAC032-like and ANAC002-like motifs. Statistical differences between *A. thaliana* and *P. patens* were assessed using Fisher’s exact tests followed by Benjamini–Hochberg correction. Asterisks indicate significant cross-species differences after FDR correction (*p<0.01, **p<0.001, ***p<0.0001).

Because the Arabidopsis β-cc detoxification pathway is strongly associated with SCL14/TGA signaling, we then examined the distribution of an as-1/TGA-like motif in relation to ATAF/NAC (ANAC002, ANAC032 and ANAC102) promoter elements (Figure 8c). The frequency of ATAF/NAC motifs was remarkably similar between species across high-response gene sets (approximately 30% of promoters of the 500 most induced genes). In contrast, the as-1-like motif showed a strong and consistent Arabidopsis bias. It was present in 49% of the top 100 Arabidopsis responders compared with 7% in Physcomitrium, in 35.5% versus 10.5% of the top 200 genes, and in 26.2% versus 13.8% of the top 500 genes. The combined as1+NAC architecture showed an even stronger separation among the most highly induced genes, occurring in 20% of the top 100 Arabidopsis genes but only 1% of the corresponding Physcomitrium set. This trend became more pronounced when absolute response amplitude was considered. Among genes induced by log_2_FC ≥ 2, as-1-like motifs were present in 53.4% of Arabidopsis promoters compared with 13.3% in Physcomitrium, whereas

ATAF/NAC motifs remained comparable between species (35.2% versus 28.3%). Among genes induced by log_2_FC ≥ 3, the as-1 frequency increased further to 72.7% in Arabidopsis but remained low in Physcomitrium (14.0%). The as1+NAC-any combination was likewise enriched in Arabidopsis at both thresholds.

Together, these analyses reveal that ATAF/NAC *cis*-regulatory signatures are broadly retained across the β-cc response in both species, whereas association of the as-1-like motif with increasing response amplitude is a distinctive feature of Arabidopsis. This suggests that evolutionary divergence of the β-cc response may involve the preferential recruitment of an as-1-centered regulatory architecture (SCL/TGA/ANAC) into the high-amplitude transcriptional response (D’Alessandro *et al*., 2018).

## 4 Discussion

Our findings reveal a divergence between β-cc responsiveness and its protective effects in *Physcomitrium patens*. Although EL induced β-cc accumulation and exogenous β-cc activated a broad stress-associated transcriptional response, β-cc failed to confer photoprotection and instead impaired photosynthetic performance.

The transcriptional response to β-cc substantially overlapped with that elicited by EL: approximately 66% of β-cc-responsive genes were also differentially expressed under EL. In addition, the rank-overlap enrichment analyses showed a strong concordance between β-cc and EL transcriptional responses, in contrast to the pattern reported in Arabidopsis. The β-cc stress-associated response included the induction of 12 of the 14 *EARLY LIGHT-INDUCIBLE PROTEIN 1* (*ELIP1*) homologs identified in Physcomitrium (Supplemental Data 1). Indeed, responsiveness to β-cc has previously been proposed to be an ancient trait, potentially present in eukaryotic predators of cyanobacteria (Calderon & Strand, 2021). Our findings are consistent with this hypothesis, while indicating that a substantial transcriptional response to β-cc does not necessarily translate into enhanced tolerance to EL. On the contrary, β-cc decreased photosynthetic performance in Physcomitrium, similarly to the responses reported in some algae and cyanobacteria, even when plants were exposed to vapor generated from only 2 μL of the compound.

Among carotenoids, the absence of comparable detrimental effects on photosynthetic performance following β-I treatment supports the specificity of the response to β-cc.

Finally, β-cc did not alter the abundance of the photosynthetic proteins examined, including Rubisco, suggesting that its effects may involve post-translational modifications or changes in the biochemical environment of the photosynthetic apparatus. In Arabidopsis, β-cc also acts primarily through signaling-mediated responses rather than direct changes in photosynthetic performance (Tiwari *et al*., 2025; Carna *et al*., 2026).

We therefore propose that the ability of Physcomitrium to respond to β-cc without acquiring increased EL tolerance reflects a divergence in the downstream detoxification machinery. The SCL14/TGAII/ANAC regulatory lineages and several associated enzyme subgroups show limited conservation relative to Arabidopsis. Consistent with this divergence, the transcriptional response to β-cc in Physcomitrium showed limited overlap with that observed in Arabidopsis. Together, these findings suggest that the recruitment of specific detoxification modules helped establish the protective effects of β-cc during plant evolution.

The distribution of the ANAC081/ANAC102-related lineage provides a particularly informative comparison. Members of this clade were recovered in terrestrial monocots but not in the aquatic monocots (*Lemna turionifera)* or *Physcomitrium patens* (Supplemental Figures S5, S6 and S7). Notably, Lemna retained candidates associated with the ANAC002-containing branches, suggesting a specific difference within the regulatory module rather than the absence of the entire β-cc-associated ATAF subgroup. The absence of the ANAC081/ANAC102-related lineage in Lemna and Physcomitrium proteomes coincides with detrimental responses to β-cc in both species, highlighting this regulatory branch as a candidate contributor to the balance between toxicity and acclimation.

This pattern raises the possibility that distinct evolutionary histories underlie the absence of ANAC081/ANAC102-like proteins in Physcomitrium and Lemna. In Physcomitrium, the absence of recognizable members of the broader β-cc-associated ATAF subgroup is consistent with diversification of this regulatory module after the bryophyte–vascular plant divergence. In Lemna, the presence of ANAC002-related proteins, together with the recovery of the ANAC081/ANAC102-related lineage in terrestrial monocots, raises the possibility of marked divergence of this branch. Indeed, the return of Lemna to an aquatic environment was accompanied by the reduction of several traits associated with terrestrial adaptation (Fang *et al*., 2025). Changes in the β-cc-associated regulatory machinery might represent another aspect of this evolutionary transition. Under this model, the contrasting physiological effects of β-cc would depend partly on the evolutionary retention and functional integration of downstream regulators.

Finally, because the molecular machinery regulating the β-cc response is conserved mainly at the level of broad protein families rather than specific regulatory lineages, we compared the *cis*-regulatory architecture of β-cc-induced genes in Arabidopsis and Physcomitrium. Several stress-associated motif families were enriched in both species. Conversely, a more focused analysis of the SCL/TGA/ANAC-associated *cis* architecture revealed a different pattern. ATAF/NAC motifs occurred at comparable frequencies among highly induced genes in the two species, whereas the as-1-like motif became progressively overrepresented with increasing response amplitude in Arabidopsis. The same trend was observed for promoters containing both as-1-like and NAC-like elements. These results suggest that the major evolutionary difference may not lie in the presence of as1- or NAC-like *cis* elements per se, but in their recruitment into an as-1-centered regulatory architecture responsive to β-cc.

A comparable separation between component conservation and signaling output has been documented for GUN1-dependent chloroplast retrograde signaling in bryophytes. GENOMES UNCOUPLED 1 (GUN1) is present in *Marchantia polymorpha* and likely originated in the streptophyte algal ancestors of land plants. MpGUN1 successfully complemented the Arabidopsis *gun1* mutant, restoring the characteristic retrograde response to inhibition of plastid translation. However, Marchantia did not exhibit the corresponding Arabidopsis-like GUN1-dependent repression of photosynthesis-associated nuclear genes (Honkanen & Small, 2022).

Together, these observations support a model in which retrograde signaling networks evolved through the recruitment and integration of ancient cellular components. Our findings suggest that responsiveness to β-cc and its capacity to promote stress tolerance are separable traits, with the latter depending on the downstream regulatory machinery. Furthermore, the distribution of the SCL14 and ANAC081/ANAC102-related lineage identifies a candidate link between the evolution of this machinery and the contrasting physiological outcomes of β-cc exposure.

## Acknowledgments

This work was funded by the European Union – NextGenerationEU, Mission 4, Component 1, CUP D53D23022160001 (IRONCROP), and CUP D53D23005110001 (LICAT).

## Author contributions: CRediT

P.K.K., C.T., M.C., E.T. and C.B. contributed to data acquisition and analysis. S.D. and A.A. supervised the work and contributed to writing the manuscript. S.D. and A.A. acquired funding.

## Data availability

Raw RNA-sequencing reads are deposited in the NCBI Sequence Read Archive under BioProject …

## Use of Artificial Intelligence

ChatGPT (OpenAI), academic account, was used as a supporting tool to improve the clarity, grammar, and overall quality of the English language used in this work. It was also used to assist with the development, revision, and debugging of Python and R scripts. All AI-generated suggestions, text revisions, and code were critically reviewed, tested, and validated by the authors.

## Supporting Information

## References

Ashburner M, Ball CA, Blake JA, Botstein D, Butler H, Cherry JM, Davis AP, Dolinski K, Dwight SS, Eppig JT, et al. 2000. Gene Ontology: tool for the unification of biology. Nature Genetics 25: 25–29.

Ashton NW, Grimsley NH, Cove DJ. 1979. Analysis of gametophytic development in the moss, Physcomitrella patens, using auxin and cytokinin resistant mutants. Planta 144: 427–435.

Avesani S, Lazazzara V, Buti M, Oberhuber M, Robatscher P, Perazzolli M. 2025. Volatile 2-Phenylethanol and β-Cyclocitral Trigger Defense-Related Transcriptional and Metabolic Changes in Grapevine Leaves Against Downy Mildew. Physiologia Plantarum 177: e70412.

Bailey TL. 2021. STREME: accurate and versatile sequence motif discovery. Bioinformatics 37: 2834–2840.

Bassi R, Pineau B, Dainese P, Marquardt J. 1993. Carotenoid-binding proteins of photosystem II. European Journal of Biochemistry 212: 297–303.

Beraldo C, Toffanin C, Morosinotto T, Alboresi A. 2025. Additive Effects of Multiple Photoprotective Mechanisms Drive Efficient Photosynthesis Under Variable Light Conditions. Plant, Cell & Environment 48: 7186–7198.

Bi G, Zhao S, Yao J, Wang H, Zhao M, Sun Y, Hou X, Haas FB, Varshney D, Prigge M, et al. 2024. Near telomere-to-telomere genome of the model plant Physcomitrium patens. Nature Plants 10: 327–343.

Britton G, Liaaen-Jensen S, Pfander H. 2012. Carotenoids: Handbook. Birkhäuser.

Burton GW, Mogg TJ, Riley WW, Nickerson JG. 2021. β-Carotene oxidation products -Function and safety. Food and Chemical Toxicology 152: 112207.

Calderon RH, Strand Å. 2021. How retrograde signaling is intertwined with the evolution of photosynthetic eukaryotes. Current Opinion in Plant Biology 63: 102093.

Carnà M, Korwin Krukowski P, Tosato E, D’Alessandro S. 2025. Apocarotenoids as Stress Signaling Molecules in Plants. Agriculture 15: 926.

Carna M, Tosato E, Krukowski PK, Parmagnani AS, Mannino G, Costa A, D’Alessandro S. 2026. β-cyclocitral elicits rapid signaling distinct from the excess-light response in Arabidopsis. : 2026.10.06.757037.

Christie WW, Han X. 2010. Lipid Analysis: Isolation, Separation, Identification and Lipidomic Analysis: Fourth Edition. Elsevier Ltd.

Cresta A, D’Alessandro S. 2023. Arabidopsis ANAC102, Chloroplastic or Nucleocytosolic Localization? Genes 14: 438.

D’Alessandro S, Ksas B, Havaux M. 2018. Decoding β-Cyclocitral-Mediated Retrograde Signaling Reveals the Role of a Detoxification Response in Plant Tolerance to Photooxidative Stress. The Plant Cell 30: 2495–2511.

D’Alessandro S, Mizokami Y, Légeret B, Havaux M. 2019. The Apocarotenoid β-Cyclocitric Acid Elicits Drought Tolerance in Plants. iScience 19: 461–473.

Dall’Osto L, Bressan M, Bassi R. 2015. Biogenesis of light harvesting proteins. Biochimica et Biophysica Acta (BBA) - Bioenergetics 1847: 861–871.

Deshpande S, Purkar V, Mitra S. 2021. β-Cyclocitral, a Master Regulator of Multiple Stress-Responsive Genes in Solanum lycopersicum L. Plants. Plants (Basel, Switzerland) 10: 2465.

Emms DM, Kelly S. 2019. OrthoFinder: phylogenetic orthology inference for comparative genomics. Genome Biology 20: 238.

Fang Y, Tian X, Jin Y, Du A, Ding Y, Liao Z, He K, Zhao Y, Guo L, Xiao Y, et al. 2025. Duckweed Evolution: from Land back to Water. Genomics, Proteomics & Bioinformatics 23: qzaf074.

Felemban A, Moreno JC, Mi J, Ali S, Sham A, AbuQamar SF, Al-Babili S. 2024. The apocarotenoid β-ionone regulates the transcriptome of Arabidopsis thaliana and increases its resistance against Botrytis cinerea. The Plant Journal 117: 541–560.

Fode B, Siemsen T, Thurow C, Weigel R, Gatz C. 2008. The Arabidopsis GRAS Protein SCL14 Interacts with Class II TGA Transcription Factors and Is Essential for the Activation of Stress-Inducible Promoters. The Plant Cell 20: 3122–3135.

Goodstein DM, Shu S, Howson R, Neupane R, Hayes RD, Fazo J, Mitros T, Dirks W, Hellsten U, Putnam N, et al. 2012. Phytozome: a comparative platform for green plant genomics. Nucleic Acids Research 40: D1178–D1186.

Grant CE, Bailey TL, Noble WS. 2011. FIMO: scanning for occurrences of a given motif. Bioinformatics 27: 1017–1018.

Gruszecki WI, Strzałka K. 2005. Carotenoids as modulators of lipid membrane physical properties. Biochimica et Biophysica Acta (BBA) - Molecular Basis of Disease 1740: 108–115.

Haslam TM, Herrfurth C, Feussner I. 2024. Diverse INOSITOL PHOSPHORYLCERAMIDE SYNTHASE mutant alleles of Physcomitrium patens offer new insight into complex sphingolipid metabolism. New Phytologist 242: 1189–1205.

Hodges DM, DeLong JM, Forney CF, Prange RK. 1999. Improving the thiobarbituric acid-reactive-substances assay for estimating lipid peroxidation in plant tissues containing anthocyanin and other interfering compounds. Planta 207: 604–611.

Honkanen S, Small I. 2022. The GENOMES UNCOUPLED1 protein has an ancient, highly conserved role but not in retrograde signalling. New Phytologist 236: 99–113.

Hunter JD. 2007. Matplotlib: A 2D Graphics Environment. Computing in Science Engineering 9: 90–95.

Jia K-P, Mi J, Ali S, Ohyanagi H, Moreno JC, Ablazov A, Balakrishna A, Berqdar L, Fiore A, Diretto G, et al. 2022. An alternative, zeaxanthin epoxidase-independent abscisic acid biosynthetic pathway in plants. Molecular Plant 15: 151–166.

Jüttner F, Watson SB, von Elert E, Köster O. 2010. β-Cyclocitral, a Grazer Defence Signal Unique to the Cyanobacterium Microcystis. Journal of Chemical Ecology 36: 1387–1397.

Kalyaanamoorthy S, Minh BQ, Wong TKF, von Haeseler A, Jermiin LS. 2017. ModelFinder: fast model selection for accurate phylogenetic estimates. Nature Methods 14: 587–589.

Kanehisa M, Goto S, Furumichi M, Tanabe M, Hirakawa M. 2010. KEGG for representation and analysis of molecular networks involving diseases and drugs. Nucleic Acids Research 38: D355–D360.

Katoh K, Standley DM. 2013. MAFFT multiple sequence alignment software version 7: improvements in performance and usability. Molecular Biology and Evolution 30: 772–780.

Landi M. 2017. Commentary to: “Improving the thiobarbituric acid-reactive-substances assay for estimating lipid peroxidation in plant tissues containing anthocyanin and other interfering compounds” by Hodges et al., Planta (1999) 207:604–611. Planta 245: 1067–1067.

Love MI, Huber W, Anders S. 2014. Moderated estimation of fold change and dispersion for RNA-seq data with DESeq2. Genome Biology 15: 550.

Lundquist PK, Davis JI, van Wijk KJ. 2012. ABC1K atypical kinases in plants; filling the organellar kinase void. Trends in plant science 17: 546–555.

Lv F, Zhou J, Zeng L, Xing D. 2015. β-cyclocitral upregulates salicylic acid signalling to enhance excess light acclimation in Arabidopsis. Journal of Experimental Botany 66: 4719–4732.

McKinney W. 2010. Data Structures for Statistical Computing in Python. SciPy 2010.

Minh BQ, Schmidt HA, Chernomor O, Schrempf D, Woodhams MD, von Haeseler A, Lanfear R. 2020. IQ-TREE 2: New Models and Efficient Methods for Phylogenetic Inference in the Genomic Era. Molecular Biology and Evolution 37: 1530–1534.

Mitra S, Estrada-Tejedor R, Volke DC, Phillips MA, Gershenzon J, Wright LP. 2021. Negative regulation of plastidial isoprenoid pathway by herbivore-induced β-cyclocitral in Arabidopsis thaliana. Proceedings of the National Academy of Sciences 118: e2008747118.

Pan N, Xu H, Chen W, Liu Z, Liu Y, Huang T, Du S, Xu S, Zheng T, Zuo Z. 2024. Cyanobacterial VOCs β-ionone and β-cyclocitral poisoning *Lemna turionifera* by triggering programmed cell death. Environmental Pollution 342: 123059.

Pinnola A, Bassi R. 2018. Molecular mechanisms involved in plant photoprotection. Biochemical Society Transactions 46: 467–482.

Ramel F, Birtic S, Ginies C, Soubigou-Taconnat L, Triantaphylidès C, Havaux M. 2012. Carotenoid oxidation products are stress signals that mediate gene responses to singlet oxygen in plants. Proceedings of the National Academy of Sciences 109: 5535–5540.

Rauluseviciute I, Riudavets-Puig R, Blanc-Mathieu R, Castro-Mondragon JA, Ferenc K, Kumar V, Lemma RB, Lucas J, Chèneby J, Baranasic D, et al. 2024. JASPAR 2024: 20th anniversary of the open-access database of transcription factor binding profiles. Nucleic Acids Research 52: D174–D182.

Rieseberg TP, Dadras A, Darienko T, Post S, Herrfurth C, Fürst-Jansen JMR, Hohnhorst N, Petroll R, Rensing SA, Pröschold T, et al. 2025. Time-resolved oxidative signal convergence across the algae–embryophyte divide. Nature Communications 16: 1780.

Roach T, Baur T, Kranner I. 2022. β-Cyclocitral Does Not Contribute to Singlet Oxygen-Signalling in Algae, but May Down-Regulate Chlorophyll Synthesis. Plants 11: 2155.

Sandmann G. 2021. Diversity and origin of carotenoid biosynthesis: its history of coevolution towards plant photosynthesis. New Phytologist 232: 479–493.

Shao N, Chen M, Xu N, Qin Y, Zhao Q, Bock R, Zhou M, Duan G, Yang J, Ji D, et al. 2026. Chloroplast sunscreening by protein condensates confers high-light tolerance. Cell 189: 5182–5197.e9.

Shumbe L, Bott R, Havaux M. 2014. Dihydroactinidiolide, a high light-induced β-carotene derivative that can regulate gene expression and photoacclimation in Arabidopsis. Molecular Plant 7: 1248–1251.

Shumbe L, D’Alessandro S, Shao N, Chevalier A, Ksas B, Bock R, Havaux M. 2017. METHYLENE BLUE SENSITIVITY 1 (MBS1) is required for acclimation of Arabidopsis to singlet oxygen and acts downstream of β-cyclocitral. Plant, Cell & Environment 40: 216–226.

Simkin AJ. 2021. Carotenoids and Apocarotenoids in Planta: Their Role in Plant Development, Contribution to the Flavour and Aroma of Fruits and Flowers, and Their Nutraceutical Benefits. Plants 10: 2321.

Sommerburg O, Langhans C-D, Arnhold J, Leichsenring M, Salerno C, Crifò C, Hoffmann GF, Debatin K-M, Siems WG. 2003. β-Carotene cleavage products after oxidation mediated by hypochlorous acid—a model for neutrophil-derived degradation. Free Radical Biology and Medicine 35: 1480–1490.

Soneson C, Love MI, Robinson MD. 2016. Differential analyses for RNA-seq: transcript-level estimates improve gene-level inferences. F1000Research 4: 1521.

Song L, Huang S-SC, Wise A, Castanon R, Nery JR, Chen H, Watanabe M, Thomas J, Bar-Joseph Z, Ecker JR. 2016. A transcription factor hierarchy defines an environmental stress response network. Science 354: aag1550.

Stephens M. 2017. False discovery rates: a new deal. Biostatistics 18: 275–294.

Stratton SP, Schaefer WH, Liebler DC. 1993. Isolation and identification of singlet oxygen oxidation products of .beta.-carotene. Chemical Research in Toxicology 6: 542–547.

Subramanian A, Tamayo P, Mootha VK, Mukherjee S, Ebert BL, Gillette MA, Paulovich A, Pomeroy SL, Golub TR, Lander ES, et al. 2005. Gene set enrichment analysis: A knowledge-based approach for interpreting genome-wide expression profiles. Proceedings of the National Academy of Sciences 102: 15545–15550.

Sun Q, Zhou M, Zuo Z. 2020. Toxic mechanism of eucalyptol and β-cyclocitral on *Chlamydomonas reinhardtii* by inducing programmed cell death. Journal of Hazardous Materials 389: 121910.

Takasaki H, Maruyama K, Takahashi F, Fujita M, Yoshida T, Nakashima K, Myouga F, Toyooka K, Yamaguchi-Shinozaki K, Shinozaki K. 2015. SNAC-As, stress-responsive NAC transcription factors, mediate ABA-inducible leaf senescence. The Plant Journal 84: 1114–1123.

Taniguchi S, Takeda A, Kiryu M, Gomi K. 2023. Jasmonic Acid-Induced β-Cyclocitral Confers Resistance to Bacterial Blight and Negatively Affects Abscisic Acid Biosynthesis in Rice. International Journal of Molecular Sciences 24: 1704.

Telfer A. 2002. What is beta-carotene doing in the photosystem II reaction centre? Philosophical Transactions of the Royal Society B: Biological Sciences 357: 1431–1470.

Thelander M, Olsson T, Ronne H. 2005. Effect of the energy supply on filamentous growth and development in Physcomitrella patens. Journal of Experimental Botany 56: 653–662.

Thimm O, Bläsing O, Gibon Y, Nagel A, Meyer S, Krüger P, Selbig J, Müller LA, Rhee SY, Stitt M. 2004. MAPMAN: a user-driven tool to display genomics data sets onto diagrams of metabolic pathways and other biological processes. The Plant Journal: For Cell and Molecular Biology 37: 914–939.

Tietz S, Hall CC, Cruz JA, Kramer DM. 2017. NPQ(T) : a chlorophyll fluorescence parameter for rapid estimation and imaging of non-photochemical quenching of excitons in photosystem-II-associated antenna complexes. Plant, Cell & Environment 40: 1243–1255.

Tiwari M, Ksas B, Légeret B, Viola S, Caffarri S, Havaux M. 2025. CYP81D11, an apocarotenoid-responsive cytochrome P450, enhances photosynthesis and stress tolerance in plants. Plant Physiology 199: kiaf476.

Tomita K, Hasegawa M, Arii S, Tsuji K, Bober B, Harada K. 2016. Characteristic oxidation behavior of β-cyclocitral from the cyanobacterium Microcystis. Environmental Science and Pollution Research 23: 11998–12006.

Vallat R. 2018. Pingouin: statistics in Python. Journal of Open Source Software 3: 1026.

Virtanen P, Gommers R, Oliphant TE, Haberland M, Reddy T, Cournapeau D, Burovski E, Peterson P, Weckesser W, Bright J, et al. 2020. SciPy 1.0: fundamental algorithms for scientific computing in Python. Nature Methods 17: 261–272.

Wang X, Cao H, Zhu Y, Zhou T, Teng F, Tao Y. 2024. β-cyclocitral induced rapid cell death of *Microcystis aeruginosa*. Environmental Pollution 348: 123824.

Wang JY, Haider I, Jamil M, Fiorilli V, Saito Y, Mi J, Baz L, Kountche BA, Jia K-P, Guo X, et al. 2019. The apocarotenoid metabolite zaxinone regulates growth and strigolactone biosynthesis in rice. Nature Communications 10: 810.

Waskom ML. 2021. seaborn: statistical data visualization. Journal of Open Source Software 6: 3021.

Wickham H. 2016. Data Analysis. In: Wickham H, ed. ggplot2: Elegant Graphics for Data Analysis. Cham: Springer International Publishing, 189–201.

Yu G. 2024. Thirteen years of clusterProfiler. The Innovation 5: 100722.

