## Supplementary figures and images for "Evolutionary insights into β-cyclocitral signaling in *Physcomitrium patens*"

### Figure_S1

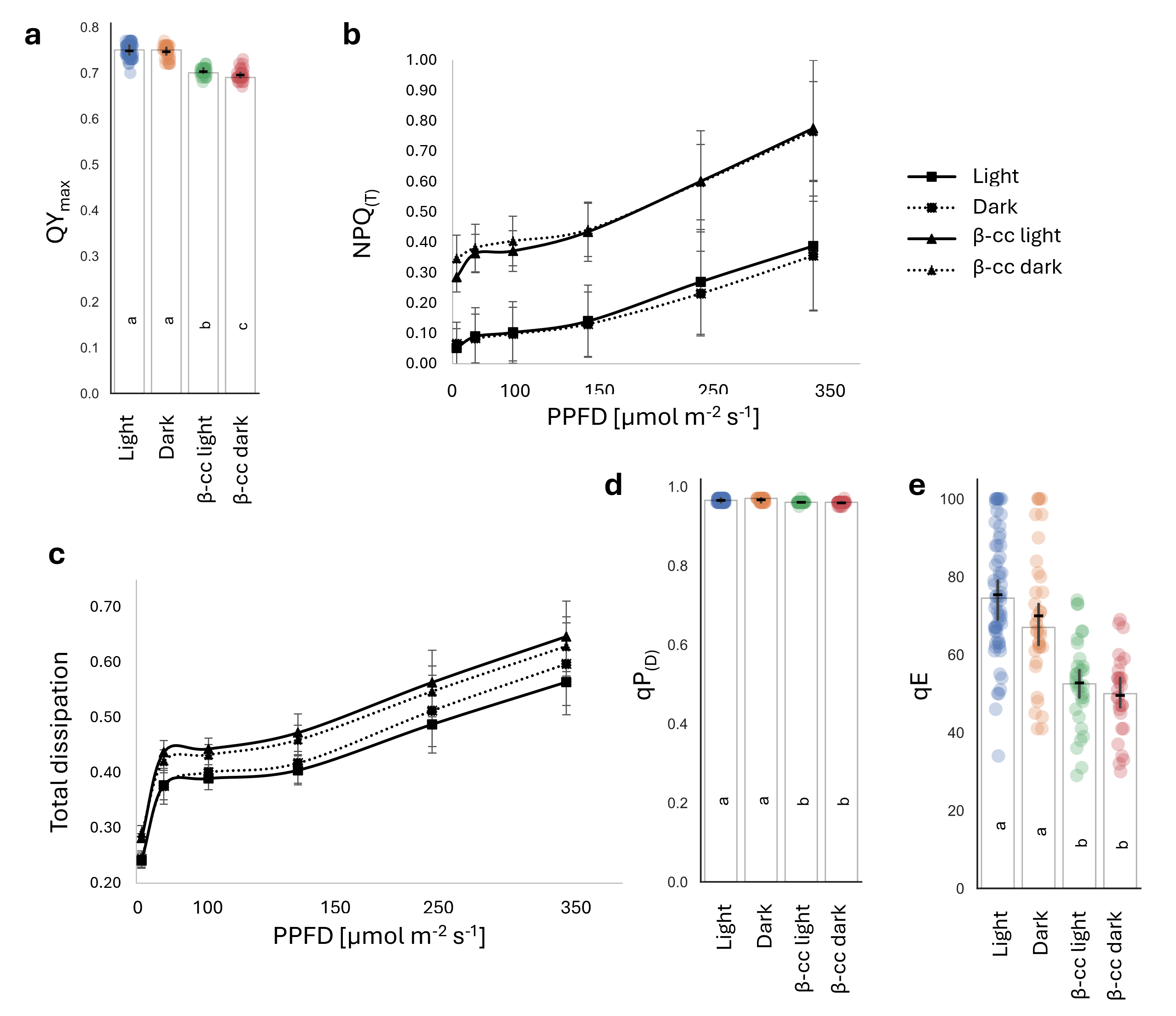

### Figure_S2

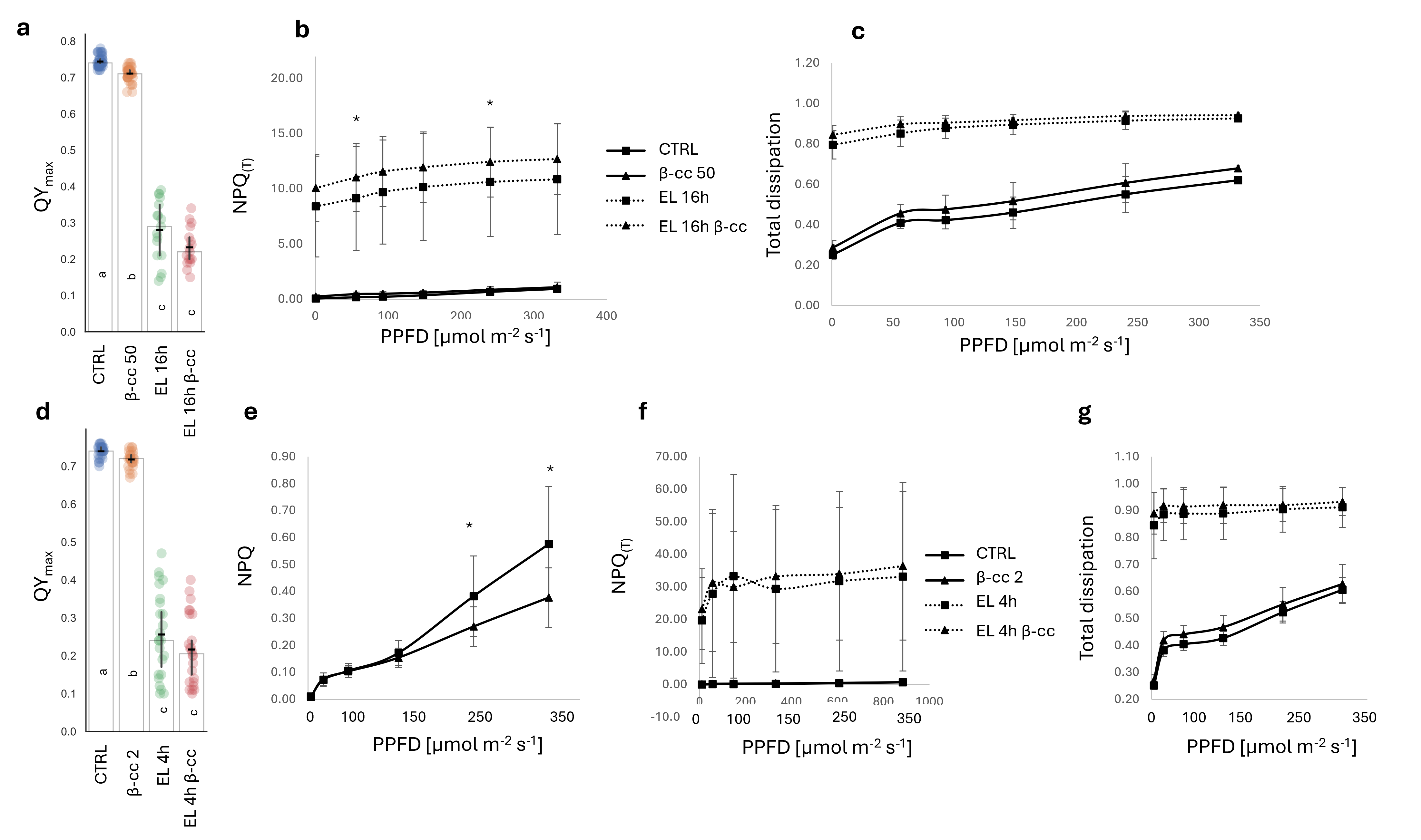

### Figure_S3

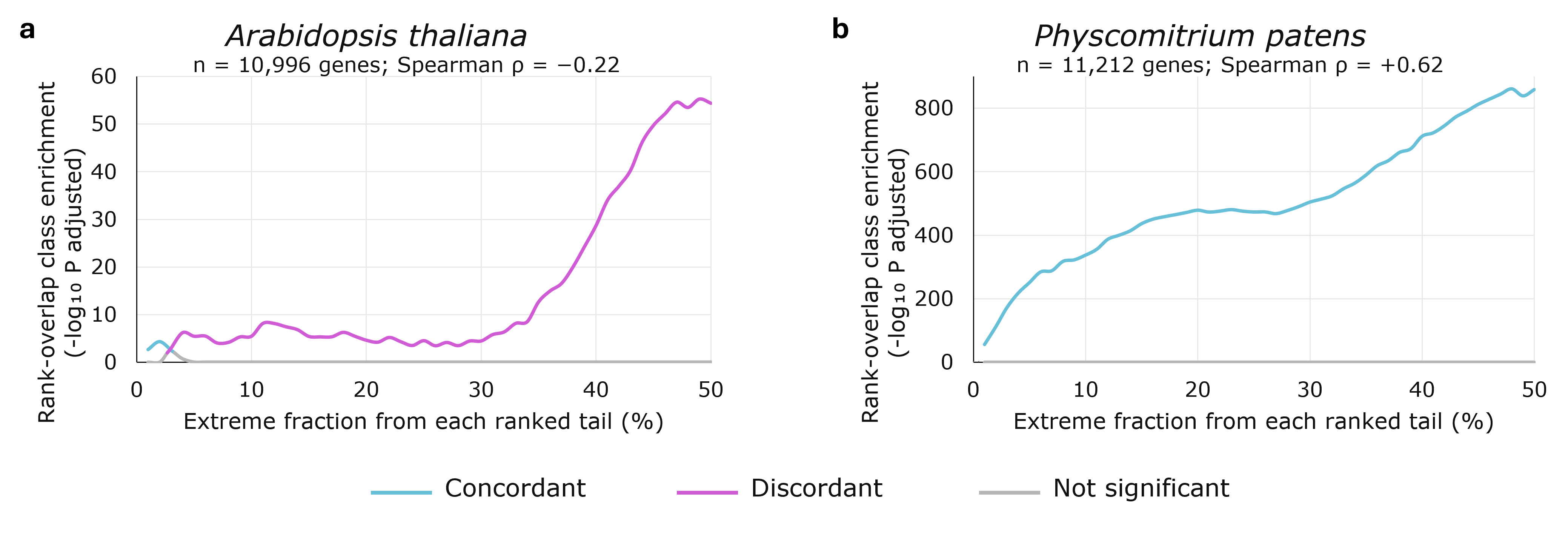

### Figure_S4

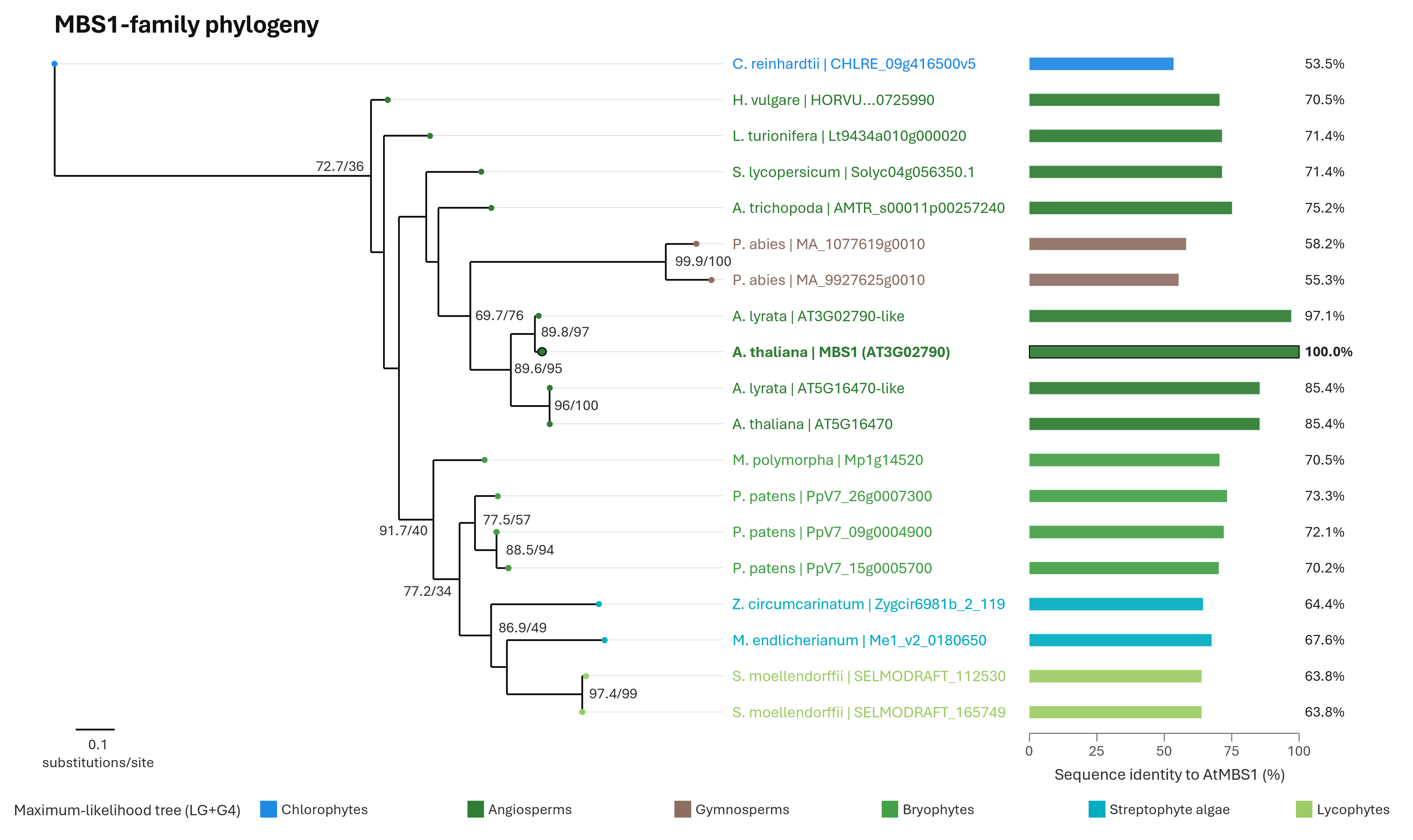

### Figure_S5

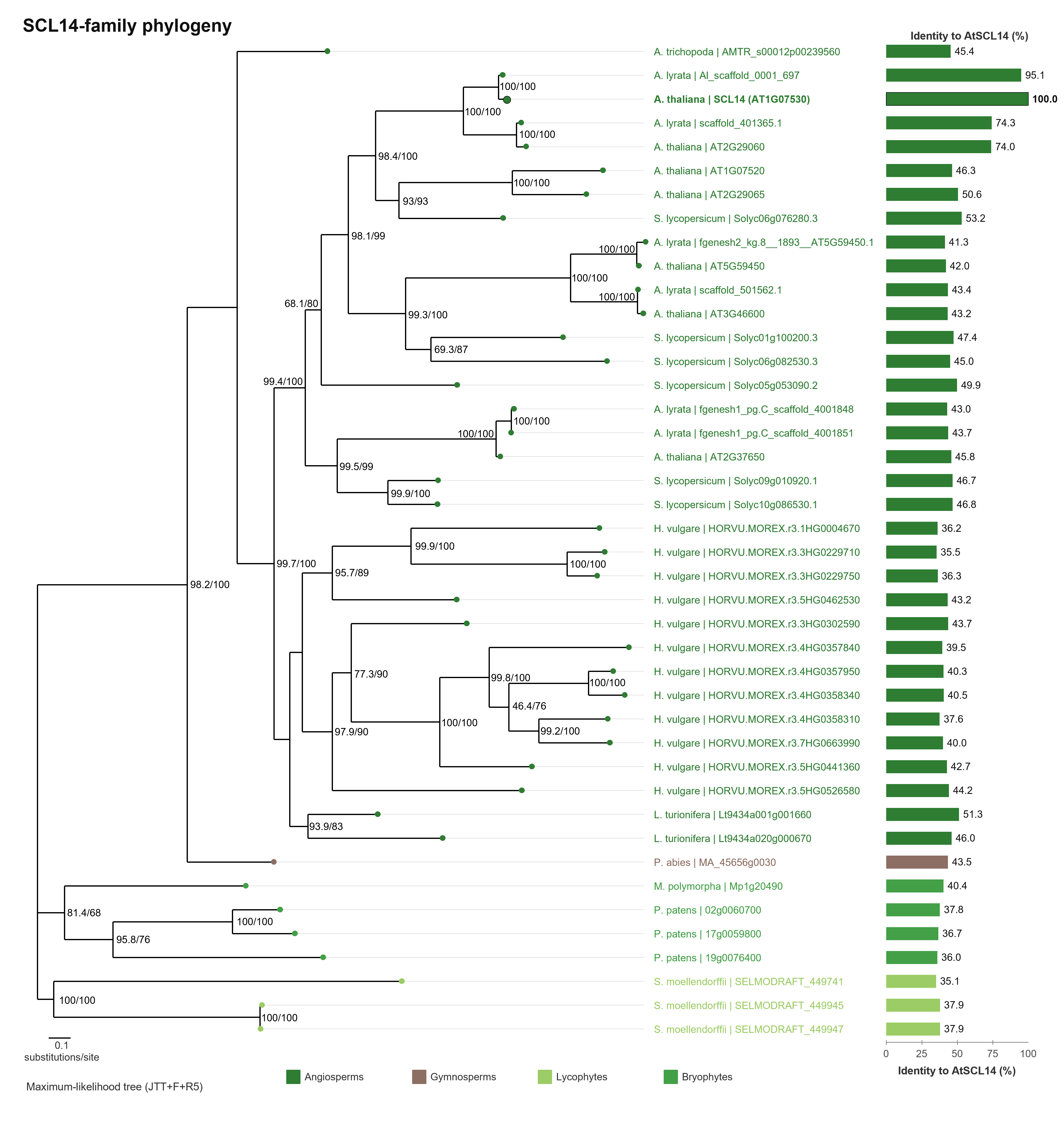

### Figure_S6

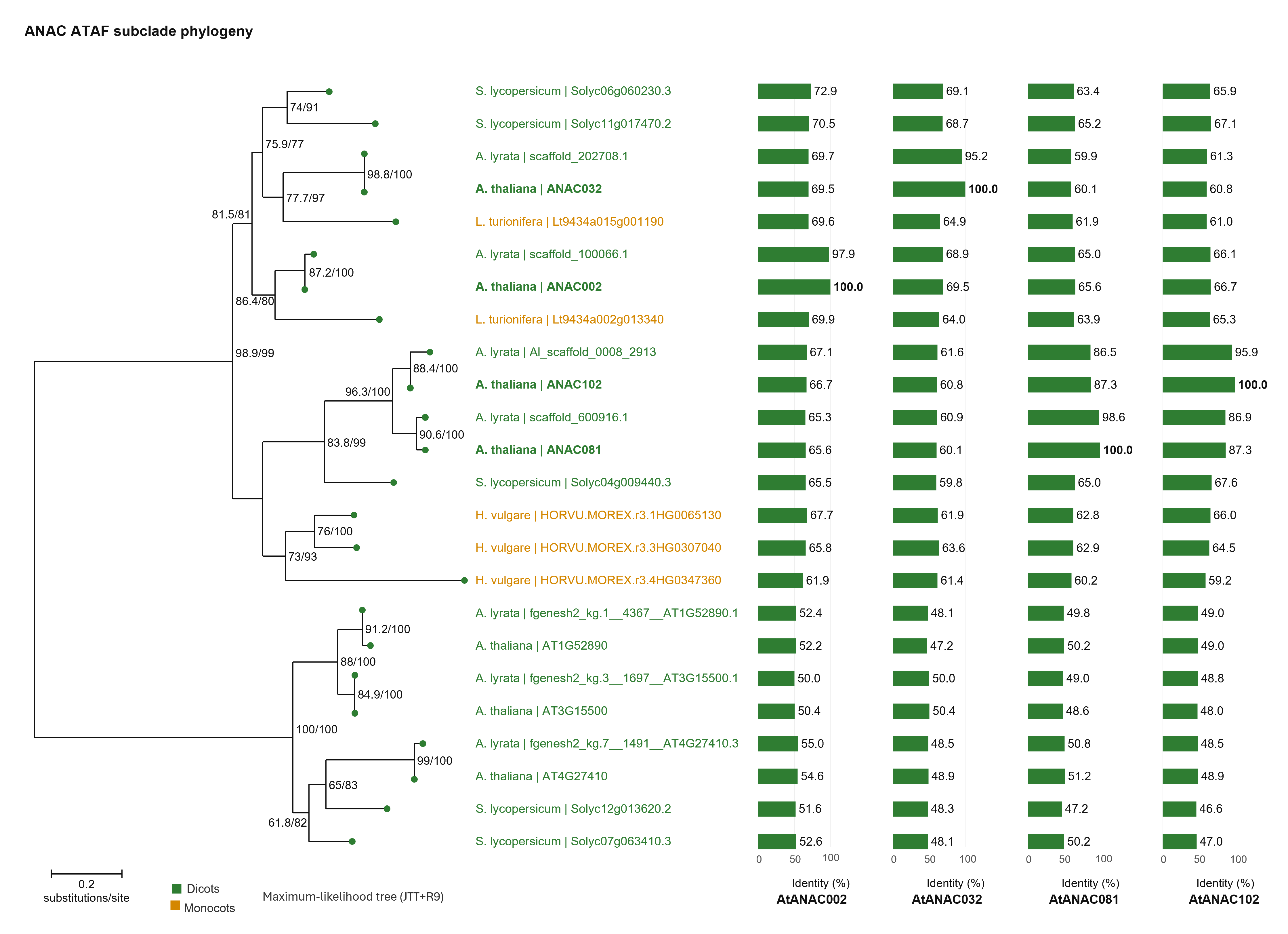

### Figure_S7

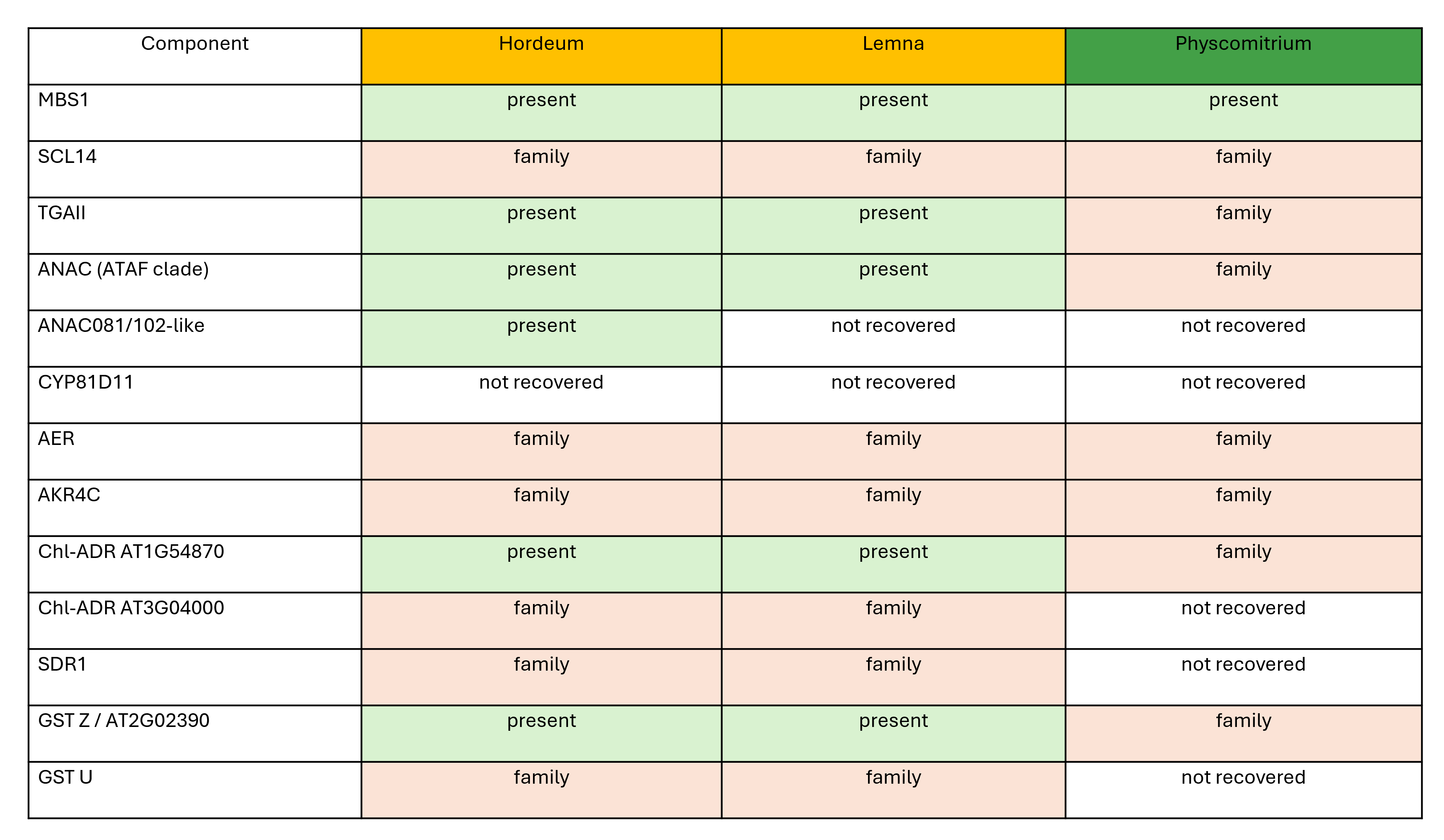

### Figure_S8

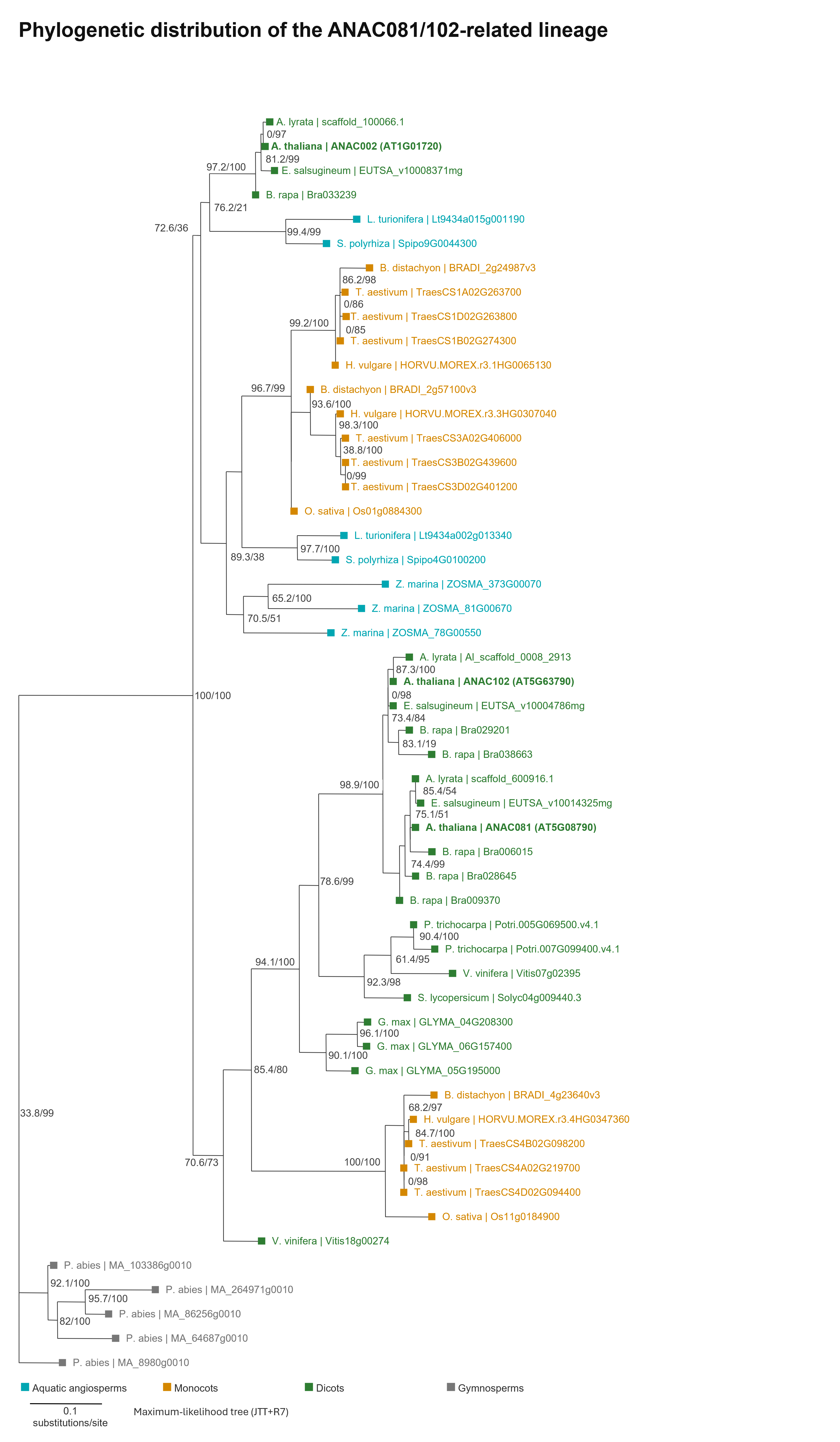
